# Leukemic stem cells hijack lineage inappropriate signalling pathways to promote their growth

**DOI:** 10.1101/2023.03.10.532081

**Authors:** Sophie G Kellaway, Sandeep Potluri, Peter Keane, Helen J. Blair, Paulynn S Chin, Anetta Ptasinska, Alice Worker, Luke Ames, Assunta Adamo, Daniel JL Coleman, Naeem Khan, Salam A. Assi, Anja Krippner-Heidenreich, Manoj Raghavan, Peter N Cockerill, Olaf Heidenreich, Constanze Bonifer

## Abstract

Acute Myeloid Leukemia (AML) is caused by multiple mutations which dysregulate growth and differentiation of myeloid cells. Cells adopt different gene regulatory networks specific to individual mutations, maintaining a rapidly proliferating blast cell population with fatal consequences for the patient if not treated. The most common treatment option is still chemotherapy which targets such cells. However, patients harbour a population of quiescent leukemic stem cells (LSCs) which can emerge from quiescence to trigger relapse after therapy. The processes that allow such cells to re- grow remain unknown. Here, we examined the well characterised t(8;21) AML sub-type as a model to address this question. Using a novel t(8;21) patient-derived xenograft model, we show that t(8;21) LSCs aberrantly activate the VEGF and IL-5 signalling pathways. Both pathways operate within a regulatory circuit consisting of the driver oncoprotein RUNX1::ETO and an AP-1/GATA2 axis allowing LSCs to re-enter the cell cycle while preserving self-renewal capacity.

## Introduction

Acute myeloid leukemia (AML) is a disease characterized by excessive production of blood progenitor cells known as blasts, which exhibit impaired differentiation capacity. This blast population is replenished by rare leukemia initiating cells, called leukemic stem cells (LSCs)^1–3^. LSCs, like healthy hematopoietic stem cells (HSCs) form a small proportion of leukemic cells and are generally quiescent^2^, they are therefore thought to be responsible for relapse following chemotherapy which targets rapidly proliferating blast cells. Thus, relapse is dependent on these cells receiving signals that induce them to re-enter the cell cycle, to proliferate to generate blasts and to repopulate the AML^4^. Whether LSCs are quiescent or proliferating is likely to be the result of transcriptional control in cooperation with signalling processes operating in the niche occupied by the cells. LSCs utilise growth control mechanisms similar but not identical to HSCs^4^, which may allow for selective targeting. For example, FoxM1 regulates the cell cycle specifically in MLL-rearranged LSCs^5, 6^. Different subtypes of AML are caused by different mutations, and we have shown that blast cells adopt subtype-specific gene regulatory networks (GRNs) maintaining the leukemic phenotype^7, 8^. It is largely unknown to what extent GRNs are already established in LSCs as compared to blast cells as the latter dominate the transcriptional signature in bulk sequencing analysis. Decoding these GRNs is further complicated by the fact that even healthy HSCs have highly diverse transcriptional profiles^9^, and LSCs also show intra and inter-patient gene expression heterogeneity^10^. Understanding the mechanisms underlying subtype specific LSC gene regulation and the transition to leukemia regeneration may reveal critical therapeutic targets driving relapse.

AML driven by the t(8;21) chromosomal translocation is one of the best characterised and common subtypes. Remission is achieved in around 90% of t(8;21) patients but they are prone to relapse associated with poor outcomes^11^. The translocation produces the RUNX1::ETO fusion protein, and the resulting AML has a unique GRN, with some aspects shared with bi-allelic *CEBPA*-mutated AMLs (biCEBPA) including a dependency on AP-1 and RUNX1 transcription factors^7, 12, 13^. The RUNX1::ETO oncoprotein is expressed under the control of the RUNX1 promoters and interferes with the normal action of RUNX1 by binding to the same sites in the genome^14, 15^. AML with t(8;21) usually carries additional mutations in signalling molecules, such as KIT or FLT3 which are thought to substantially contribute to full leukemic transformation^16, 17^. Furthermore, both cell extrinsic and intrinsic signalling are known to play roles in t(8;21) growth, whereby activation of the AP-1 pathway upregulates transcription of signalling and cell cycle genes^18–20^. t(8;21) AML is therefore an attractive model to study LSC activation and to identify targets aimed at preventing relapse. In this study, we determined the genome-wide t(8;21) LSC-specific open chromatin structure and gene expression profile. We also profiled LSCs at the single cell level using single cell RNA-Seq (scRNA-Seq) which, together with perturbation experiments using a novel PDX model for t(8;21) AML, identified the growth factors VEGFA and IL-5 and their receptors as key factors aberrantly driving the growth of this specific LSC subtype. Furthermore, we identify an oncoprotein driven transcription/signalling circuit, dependent upon the AP-1 family of transcription factors as mediators of VEGF/IL-5 signalling that regulates the balance between LSC maintenance and blast growth.

## Results

### t(8;21) LSCs exhibit mutation-specific gene expression and chromatin accessibility profiles

In this study we employed t(8;21) AML as an archetypal model system to gain an understanding of the factors which activate LSC growth and drive relapse following chemotherapy. The development of t(8;21) AML from pre-leukemic cells is typically driven by mutations in signalling molecules such as KIT. To examine whether there is a mutation subtype-specific or global mechanism underlying the control of the LSC growth status, we defined bulk transcriptional signatures for LSCs and blasts purified from two t(8;21) bone marrow samples (referred to from hereon as t(8;21) #1 and #2). Mutation profiling of bulk AML cells revealed that sample #1 had mutations in several key hematopoietic regulator genes including *GATA1*, *KIT* and *WT1* at allele frequencies of 40% and higher, while sample #2 carried two FLT3 internal tandem duplications (ITD) and a *RAD21* indel at allele frequencies ≤ 10% (Table 1). Cells were sorted for the CD34+/CD38- surface marker pattern to enrich for LSCs, with the CD34+/CD38+ fraction comprising leukemic blasts (Figure S1A)^2, 21^, followed by genome-wide profiling of gene expression RNA-seq and open chromatin regions by ATAC-seq (Figure 1A). We used colony forming assays to verify the sorted populations, with 4 colonies per 1000 cells observed from sorted LSCs and zero colonies from blasts from t(8;21) #2 (Figure S1B). QRT-PCR with specific primers targeting the fusion confirmed that these colonies expressed RUNX1::ETO and that they were not generated from contaminating wild-type cells (Figure S1C). Cells from t(8;21) #1 did not form colonies, but this is commonly observed for t(8;21) primary samples.

**Figure 1:**
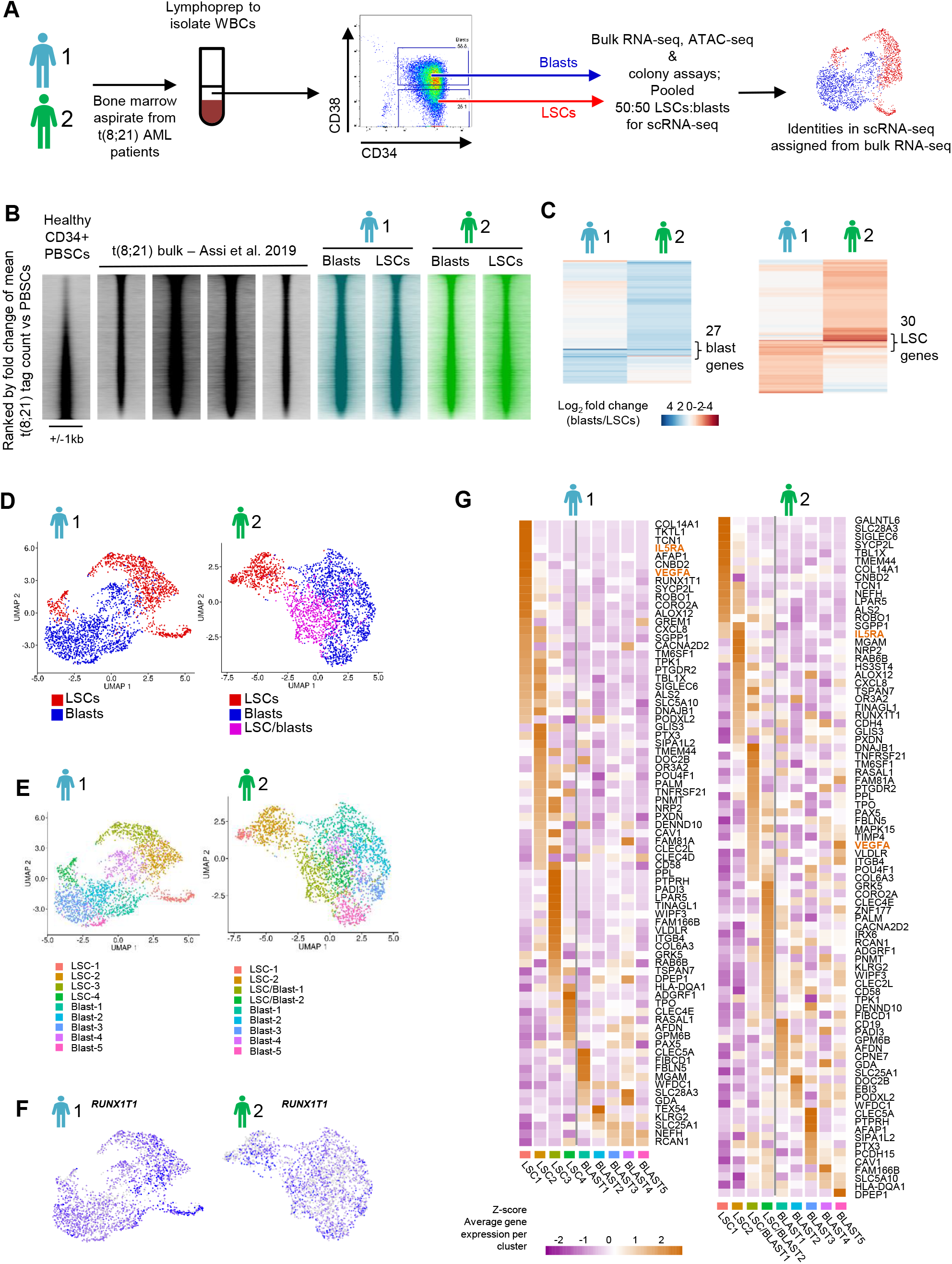
Subtype specific gene expression and chromatin accessibility is established in LSCs. **(A)** Schematic showing how patient bone marrow cells were sorted into LSCs and blasts for bulk RNA-seq, ATAC-seq, single cell RNA-seq and colony forming assays. **(B)** DNaseI-seq in t(8;21) patients and healthy CD34+ PBSCs^7^ was ranked by the fold change of the average tag count in distal peaks and represented as density plots (+/-1kb of the summit). ATAC-seq on sorted LSCs and blasts was plotted along the same axis. **(C)** Heatmaps showing the log_2_ fold change expression (blasts vs LSCs) of the genes defined in (B) as differential in both patients, with the core set of concordantly 2-fold differential genes indicated. **(D)** UMAP plots of scRNA-seq for both patients, where blue dots indicate cells assigned as blasts and red dots indicate cells assigned as LSCs. Purple dots on the second patient indicate intermediate type cells which could not confidently be assigned as blasts or LSCs. **(E)** Cell subclusters identified in each patient projected onto the UMAP plot. **(F)** Expression of *RUNX1T1* projected onto the UMAP plot, where blue indicates the normalised UMI count. **(G)** Heatmaps with hierarchical clustering showing Z-scores of average gene expression per cluster of t(8;21) specific genes.

**Table 1:** Details of mutations in patient AML cells additional to the t(8;21) translocation, obtained from West Midlands Regional Genetics Laboratory.

| Patient |  | Gene | Mutation |  | VAF |
| --- | --- | --- | --- | --- | --- |
| **t(8;21) #1** | Relapse | ETV6 | c.313_314insGG | NP_001978 p.R105fs | 45% |
|  |  | GATA1 | c.158C>A | NP_002040 p.A53D | 51% |
|  |  | KIT | c.1253_1255del | NP_001087241<br>p.418_419del | 61% |
|  |  | NOTCH1 | c.4898G>A | NP_060087 p.R1633H | 54% |
|  |  | NOTCH1 | c.6980G>A | NP_060087 p.R2327Q | 48% |
|  |  | WT1 | c.420_421insGTGTGCGA | NP_001185481<br>p.R141fs | 39% |
| **t(8;21) #2** | Presentation | RAD21 | c.1645delinsGGGGGTACT | NP_006256<br>p.Q549Gfs*66 | 10% |
|  |  | FLT3 | Internal Tandem Duplication |  | 3% |
|  |  | FLT3 | Internal Tandem Duplication |  | 1% |
| **t(8;21) #3** | Relapse | Unknown |  |  |  |
| **t(8;21) #4** | Relapse | KIT | c.2435A>T | NP_000213.1 p.D816V | 48% |
|  |  | TET2 | c.4179delA | NP_001120680<br>p.T1393fs | 94% |

In a previous study we used DNaseI-seq to profile open chromatin in CD34+ cells from four patients to show that t(8;21) AML adopts a reproducible subtype-specific chromatin accessibility pattern^7^. We compared these DNaseI-seq data from bulk CD34+ AML cells with ATAC-seq data derived from CD34/CD38-sorted t(8;21) LSCs and blasts, and with healthy CD34+ PBSCs. This experiment revealed that the t(8;21)-specific open chromatin signature, as compared to healthy cells was also found in LSCs (Figure 1B). However, when compared directly to blasts and ranked by the fold-change between the two, the LSC chromatin accessibility profile was not identical to that of blast cells (Figure S1D). Beyond the general t(8;21) signature, the open chromatin sites specific for LSCs were enriched for GATA motifs (Figure S1D and S1E) whilst the blast specific sites were enriched for C/EBP and PU.1 motifs, indicating a more mature myeloid epigenomic landscape in the blasts and a more immature accessibility pattern in LSCs^22, 23^. RNA-seq data from sorted cells was used to identify blast and LSC enriched genes for each patient (Figure S1F). These genes were then compared between the two patients, and despite diverse genetic backgrounds beyond the t(8;21) a shared signature of 27 blast genes and 30 LSC genes could be identified (Figure 1C, Table S1). Blast-specific genes included *CD38* and the *AZU1*, *PRTN3, ELANE* and *CFD* mature neutrophil granule protein gene cluster, plus *MPO* and *LYZ.* LSC-specific genes included *KLF9, DUSP5, RHOC* and four beta globin genes. The latter are known to be active in multipotent hematopoietic progenitor cells^24^. The majority of the remaining LSC and blast genes specific to each patient were either not specific to one population or, whilst showing the same trend, did not reach the 2-fold difference threshold. LSC-specific genes were strongly associated with LSC-specific accessible chromatin whilst for blast specific genes the correlation was weaker (Figure S1D). This finding may indicate that blast-gene cis-regulatory elements are already accessible and primed in LSCs.

We next performed scRNA-seq on sorted LSCs and blasts because bulk RNA-seq neither reveals the heterogeneity present within the LSC population, nor how they transit to blasts. Purified LSCs were pooled with blasts prior to sequencing to better capture this rare population of interest. Cells were assigned *in silico* as LSC or blasts (Figure 1D) based on the bulk RNA-seq data. Patient 2 bone marrow contained an intermediate population of LSCs which had already begun to express blast genes in addition to LSC genes. Several clusters were identified within the scRNA-seq – 5 blast populations and 4 LSC populations, with two of the LSC populations in patient 2 assigned as a transitional population (LSC/blast; Figure 1E). Expression of the *RUNX1T1* (ETO) transcript, which is not normally expressed in healthy cells^25^, was detected in 1825/2489 cells in patient 1 and 1480/2546 cells in patient 2, across all clusters confirming that they were all AML or pre-leukemic (Figure 1F). Clusters showed good purity as defined by specific cluster markers (Figure S1G). We then examined which clusters expressed AML-specific genes. This analysis defined 88 genes whose expression was at least 2-fold higher on average in t(8;21) patients compared to other AML subtypes or healthy CD34+ PBSCs (Figure S1H) and plotted the Z-score of the expression of these 88 genes across the clusters (Figure 1G). The majority of these genes were most highly expressed in the LSC clusters, including genes known to be important for the t(8;21) phenotype such as *POU4F1* and *PAX5*^26, 27^. Together these data show that the t(8;21)-specific regulatory network found in blasts is also found in LSCs.

### LSC and blast cells show cell cycle-specific gene expression heterogeneity

We next sought to identify specific genes regulating the growth status of LSCs and blast cells. We first assigned a cell cycle status to each cell using scRNA-seq data and allocated the cells back to their clusters. This analysis demonstrated a significant intra-patient heterogeneity since we found a higher proportion of G_0_/G_1_ cells in patient 1 LSCs compared to blasts, but similar proportions of G_0_/G_1_ cells in patient 2 LSCs and blasts (Figures 2A and S2A). Genes specifically expressed in cells from each cell cycle phase were identified, including and beyond those used to assign the cell cycle stage. Whilst S and G_2_/M phase LSCs and blasts share similar gene expression profiles and are dominated by factors essential for cell cycle regulation, we found significant differences in G_0_/G_1_ gene expression between LSCs and blasts (Figure S2B). This analysis also revealed significant heterogeneity in patient 2 in cells belonging to each phase except for G_2_/M. Clinical data revealed that this patient suffered from an infection and carried a RAD21 mutation which affects the cohesin complex and may have perturbed the proportion of cells entering and leaving the cell cycle. However, when we directly compared the gene expression pattern in G_0_/G_1_ LSCs and G_0_/G_1_ blasts from the two patients, we found that similar genes were expressed specifically in LSCs or blasts. G_0_/G_1_ LSCs expressed genes primarily associated with transcriptional control and negative regulation of the cell cycle, whilst the G_0_/G_1_ blast specific expression pattern was dominated by genes associated with translation and telomere maintenance, including elongation factors and ribosomal protein genes (Figure 2B, Table S2). The difference in expression of translation factors is reminiscent of the control of protein synthesis rates via phosphorylation of 4E-BP1 by which healthy HSCs regulate growth and quiescence^28^. Our data suggest that LSCs may use a related mechanism to control growth via signalling responsive control of translation. To investigate whether this was the case we performed mass cytometry on cultured cells from patient #2 and a further t(8;21) patient #3. Proliferation, as determined by Ki67 was higher in CD34+/CD38+ blasts than in CD34+/CD38- LSCs in both patients as expected (Figure 2C). Phosphorylation of AP-1 associated proteins CREB, JUN and JNK1/JNK2 was high in both patients, and more so in blasts than LSCs. Furthermore, blasts contained increased levels of phosphorylated 4- EBP1 and S6 which are directly involved in control of protein translation (Figures 2C and S2C). In comparison, the STAT pathways and NF-κB were not differentially active. Together these data show that concordant with their quiescence, LSCs display reduced signalling influencing translation and the AP-1 pathway.

**Figure 2:**
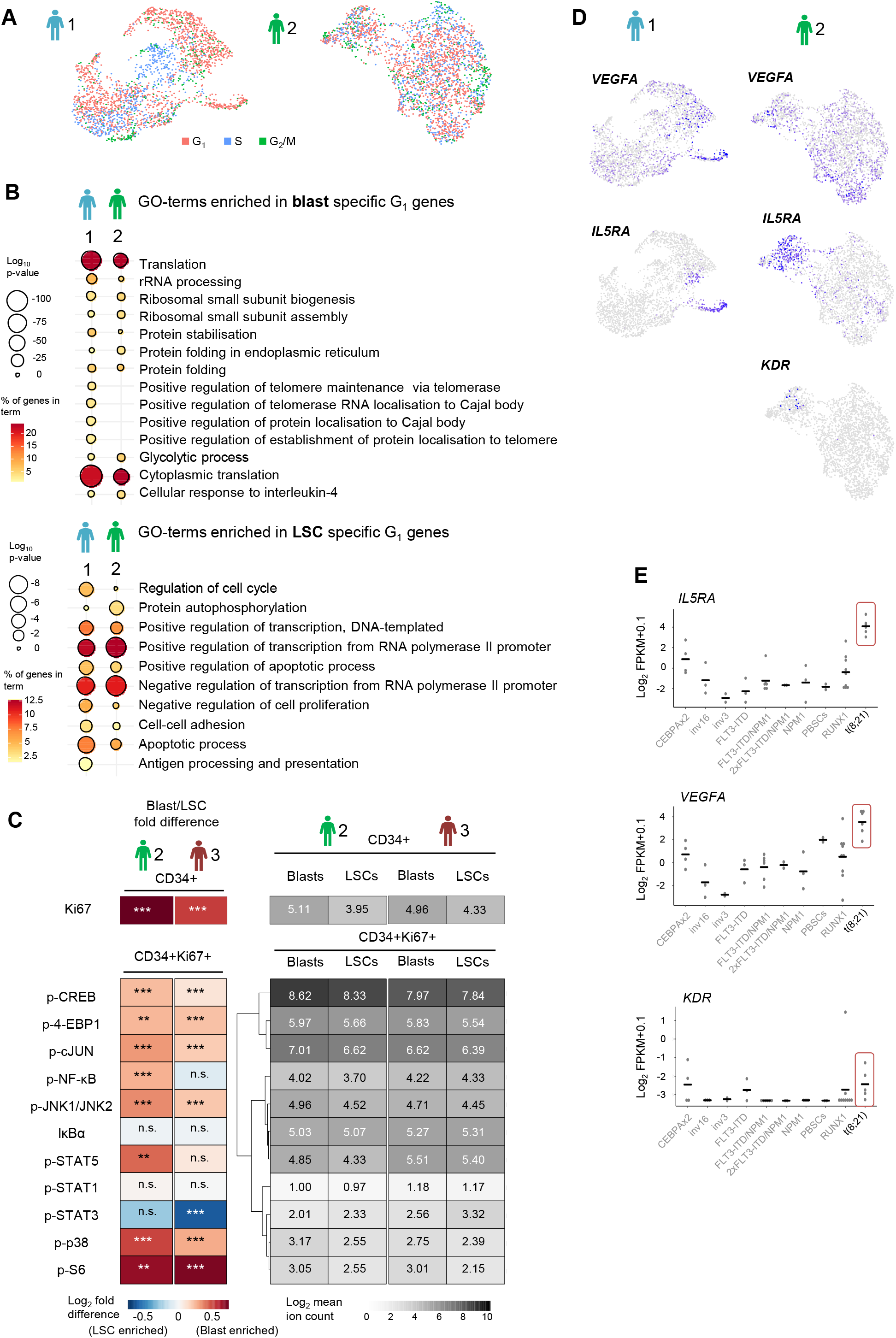
t(8;21) AML LSCs are differentially signalling responsive. **(A)** The assigned cell cycle stage of each cell projected on to the UMAP plots **(B)** Bubble plot showing enriched GO-terms generated from blast- or LSC-specific genes from G_1_ cells only, the colour scale indicates the % of genes in the GO-term which were found in the specific gene list, and the size of the bubble indicates the log_10_ p-value of the enrichment of the term. **(C)** Heatmaps showing the log_2_ fold difference between mean ion counts in blasts and LSCs (left) and the log_2_ mean ion count (right) from mass cytometry on two patients. Ki67 is shown from total CD34+ cells, all other markers are shown from CD34+Ki67+ cells. P-values for blast/LSC differences are indicated by n.s. > 0.001, * < 0.001, ** < 1e-^5^, *** < 1e-^10^ using Student’s T-tests. Patient 2 LSC n=414, blasts n=4486, patient 3 LSC n=6236, blasts n=4229. **(D)** Expression of *VEGFA, IL5RA* and *KDR* projected onto the UMAP plots, where blue indicates the normalised UMI count. **(E)** Normalised log_2_ FPKM of *IL5RA, VEGFA* and *KDR* in AML with different driver mutations and healthy CD34+ PBSCs^7^. Horizontal bars indicate the median of all samples.

**Table 2:** Antibody titres for CyTOF CyTOF experimental workflow

| **Metal** | **Markers** | **Vol (µl) / test** |
| --- | --- | --- |
| 89Y | CD45 | 1.0 |
| 106Cd | Barcode | 0.75 |
| 110Cd | Barcode | 0.75 |
| 111Cd | Barcode | 0.75 |
| 112Cd | Barcode | 0.75 |
| 113Cd | Barcode | 0.75 |
| 114Cd | Barcode | 0.75 |
| 115In | Barcode | 0.75 |
| 116Cd | Barcode | 0.75 |
| 148Nd | CD34 | 0.4 |
| 149Sm | p4E-BP1 | 0.75 |
| 150Nd | pSTAT5 | 0.5 |
| 153Eu | pSTAT1 | 0.5 |
| 156Gd | p38 | 0.5 |
| 158Gd | pSTAT3 | 0.5 |
| 159Tb | p-cJun | 1 |
| 164Dy | IkBalpha | 0.5 |
| 165Ho | CD117 | 0.75 |
| 166Er | NFkB.p65 | 0.6 |
| 167Er | CD38 | 0.5 |
| 172Yb | ki67 | 0.75 |
| 173Yb | p-Jnk1/Jnk2 | 1 |
| 175Lu | pS6 | 0.5 |
| 176Yb | pCREB | 0.4 |
| 198Pt | Barcode | 0.75 |
| 103Rh | DNA | 500 µM |
| 194Pt | LIVE/DEAD |  |

### Aberrant VEGF and IL-5 signalling in t(8;21) AML drives LSC activation and promotes the growth of a novel serially transplantable t(8;21) PDX model

To identify candidate genes which may control LSC growth, we screened for genes associated with cell signalling processes which were specific to t(8;21) LSCs. Signalling mutations such as in the KIT gene giving rise to constitutively active receptor molecules appear to be insufficient to initiate LSC growth despite being found equally in LSCs and blasts. This was also the case here, with t(8;21) #1 harboring a KIT mutation and quiescent LSCs (Figure S2A)*VEGFA* and *IL5RA mRNAs* were found to be both largely t(8;21) specific and strongly enriched in LSCs (Figures 1G, 2D and 2E). Furthermore, the *VEGFA* receptor *KDR* was also aberrantly up-regulated in t(8;21) AML as compared to healthy PBSCs and all other AML subtypes except for biCEBPA (Figure 2E). In patient 2 we could detect single LSCs expressing the VEGFA receptor *KDR,* albeit at a low level (Figure 2D) and in patient 1 *KDR* transcripts were detected in the bulk RNA-seq (raw FPKM in LSCs: 0.23, in blasts < 0.01). *VEGFA* was expressed in some blasts, particularly in patient 2 and was generally not co-expressed with *KDR* (Figures 2D and S2D). *GATA2* showed a high degree of co-expression with *VEGFA*, *KDR* and *IL5RA* (Figure S2E-F). Pseudotime trajectory analysis confirmed that these *GATA2* high, *IL5RA/KDR/VEGFA* positive LSCs were at the apex of the differentiation hierarchy (Figure S2F-G).

To assess the roles of IL-5 and VEGF signalling we used two t(8;21) cell line models: Kasumi-1 and SKNO-1. IL-5 signalling was not assessed in Kasumi-1 as this cell line does not express *IL5RA* nor display accessible chromatin at this locus^29^. We cultured cell lines in the presence of exogenous VEGF or IL-5 to maximally stimulate their respective receptors. In all cases the growth rate increased after addition of the cytokines (Figures 3A-D). We next used the VEGFA inhibitor bevacizumab^30^, with no additional VEGF (as the AML cells express it already), and the IL5RA inhibitor benralizumab^31^ with or without exogenous IL-5, to test whether the inhibitors would abrogate growth stimulation. Both inhibitors reduced growth rates (Figures 3A-C and S3A-C), although growth was not fully blocked which was expected as not all cells express IL5RA or KDR on the surface (Figure S3D). Response to benralizumab was more pronounced with the addition of exogenous IL-5 (Figures 3B and S3B). Addition of IL-5 could not compensate for the dependency of SKNO-1 on GM-CSF, even though these cytokines signal via the same receptor beta chain (Figure S3E). Our data therefore show that VEGF and IL-5 signalling promote the growth of t(8;21) AML cells.

**Figure 3:**
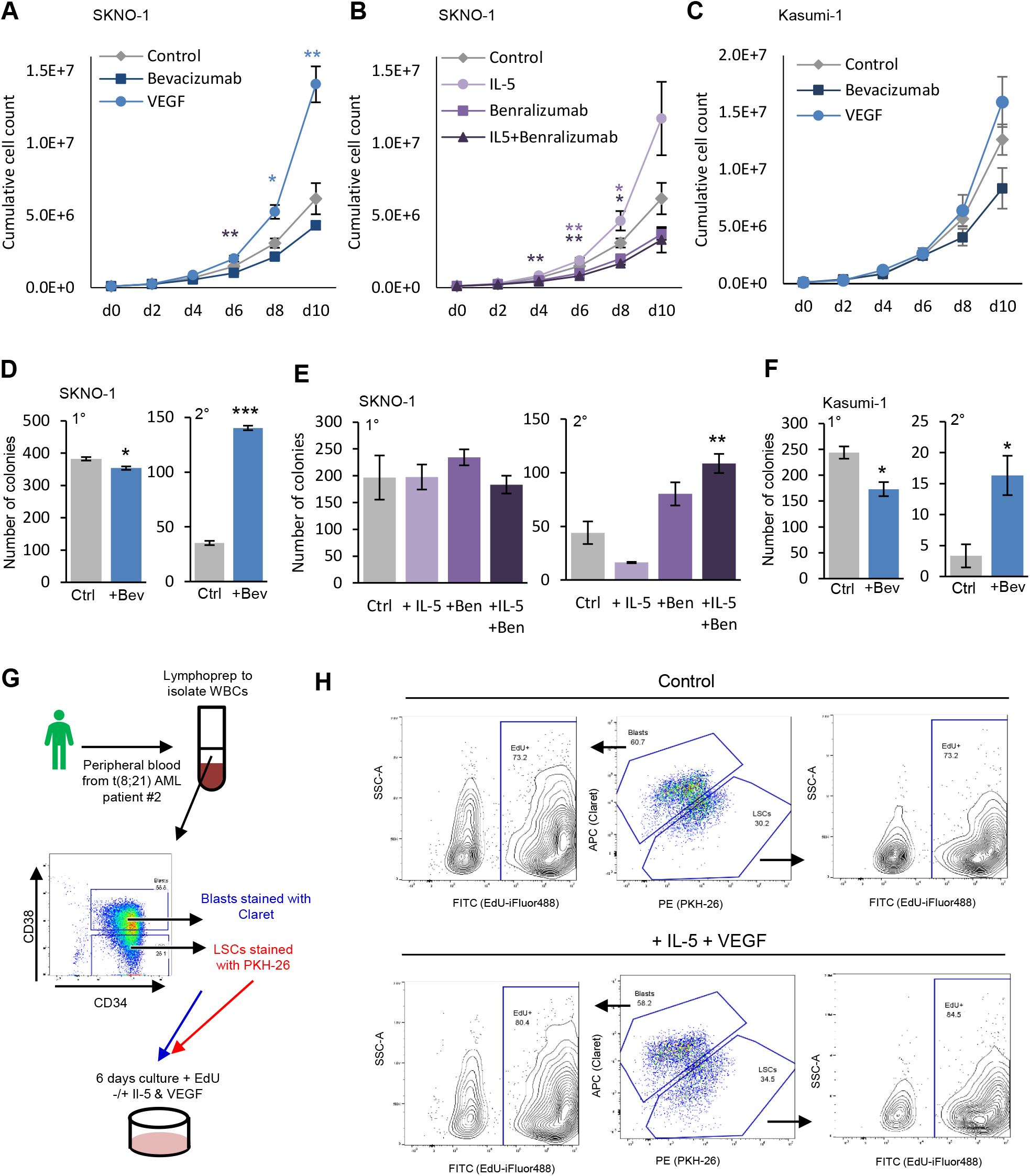
Aberrant VEGF and IL-5 signalling in t(8;21) AML drives LSC growth (A-C) SKNO-1 **(A-B)** and Kasumi-1 cells **(C)** were grown with bevacizumab, VEGF or media alone control **(A,C)** or IL-5, benralizumab, or IL-5 and benralizumab **(B)** for 10 days, with mean counts every two days indicated by the points, error bars indicate SEM. Controls are shared in A and B. n= 3-6 for each condition. **(D-F)** Primary (1°) and secondary (2°) replating colony forming assays +/- bevacizumab with SKNO-1 **(D)** and Kasumi-1 **(F)** and with IL-5, benralizumab or both in SKNO-1 **(E)**, bars indicate the mean of 3 replicates and the error bars indicate SEM**.(G)** Schematic showing how the LSC proliferation assay was conducted. **(H)** Flow cytometry plots identifying LSCs (stained with PKH-26, detected in the PE channel) and Blasts (stained with Claret, detected in the APC channel), with EdU (stained with iFluor488 and detected in the FITC channel) measured in each population separately. * indicates p <0.05, ** p<0.01, *** p<0.005 using Student’s T-tests **(D-F)** or two-way ANOVA with Dunnett correction for multiple comparisons at each time point, shown for growth curves in the same colour as the treatment group is plotted **(A-C)**.

We then carried out colony forming assays in the presence of the VEGF and IL-5 inhibitors bevacizumab and benralizumab (+/- additional IL-5). In both cell lines bevacizumab led to a small reduction in the number of colonies formed initially in concordance with the reduced growth rate, whilst benralizumab had no impact upon primary colony forming capacity (Figures 3D-F). However, when the colonies were replated a significantly higher number of colonies were formed in the presence of bevacizumab or benralizumab+IL-5 indicating a higher capacity for self-renewal (Figures 3D-F). Thus, blocking VEGF or IL-5 signalling stalls the cells in an increased self-renewing state.

Our next experiment directly studied whether IL-5 and/or VEGF signalling were indeed capable of stimulating the growth of primary LSCs. To this end, we used different membrane tracking dyes to separately label purified LSCs and blasts from t(8;21) patient #2 peripheral blood grown in cytokine-rich media with or without IL-5 and VEGF (Figures 3G and 3H). Similar proportions of LSCs and blasts were detected at the end of each assay with or without IL-5/VEGF, comparable to the proportion which were sorted and stained at the start, confirming the reliability of the membrane stains (Figure 3H). The fidelity of the gates was also confirmed by staining known proportions of unsorted cells. Both LSCs and blasts proliferated during the experiment in response to the cytokines present in the basic culture medium which includes IL-3 and GM-CSF. After the addition of IL-5 and VEGF, proliferation as measured by EdU incorporation increased, from 73% to 80% in the blasts and from 73% to 85% in LSCs. Results were the same regardless of which dye was used for which cell population (Figure S3F). Notably, the membrane dye was more variably detected with addition of IL- 5 and VEGF - particularly for the LSCs - due to dilution following cell division.

In order to generate an unlimited source of human primary t(8;21) cells we developed a patient-derived xenograft (PDX) generated from t(8;21) patient #4 who relapsed with a KIT D816V mutation. This is - to our knowledge - the first PDX from a t(8;21) patient capable of serial re-engraftment^32^, which can be cultured *ex vivo* but does not form colonies. Upon secondary engraftment the PDX maintained the gene expression pattern of the original patient cells (Figure S4A) with a leukaemia initiating cell frequency of >10^-3^ (Figure S4B). As with the cell lines, addition of VEGF or IL-5 to cultured PDX cells stimulated growth, whilst the inhibitors both reduced growth (Figures 4A-C). Combining VEGF and IL-5 in the PDX culture did not have an additive growth effect (Figure 4A). Healthy CD34+ cells showed no response to bevacizumab or benralizumab in the effective dose range observed in Figure 4 for the t(8;21) cells (Figures S4C-D). We then tested inhibition of VEGFA and IL5RA *in vivo* by injecting PDX cells intra-femorally into NSG mice and treating the animals for 41 days with bevacizumab or benralizumab (Figure 4D). Engraftment was measured by sampling peripheral blood after 72, 84 and 92 days, and bone marrow was taken at the endpoint on day 92 from the injected (right) femur and the contra-lateral (left) leg. Fewer human CD45+ cells were found in peripheral blood samples from treated mice compared to vehicle only controls (Figures 4E-F and S4E). All hCD45+ cells measured in peripheral blood were CD34+ and CD33+ showing that the cells underwent little or no myeloid differentiation. KDR and IL5RA positive cells were found predominantly in the LSC compartment of the recovered PDX cells (Figures 4G-H), demonstrating that the PDX model faithfully recapitulates the phenotype of the primary cells from the two patients. These KDR/IL5RA double positive LSCs were depleted by both inhibitors, with KDR positive LSCs further depleted by bevacizumab only (Figures 4I-J and S4F). We also noted a modest increase in the proportion of CD34-/CD11b+ cells indicative of more mature cells in the non-injected bone marrow only with treatment (Figures S4G-H).

**Figure 4:**
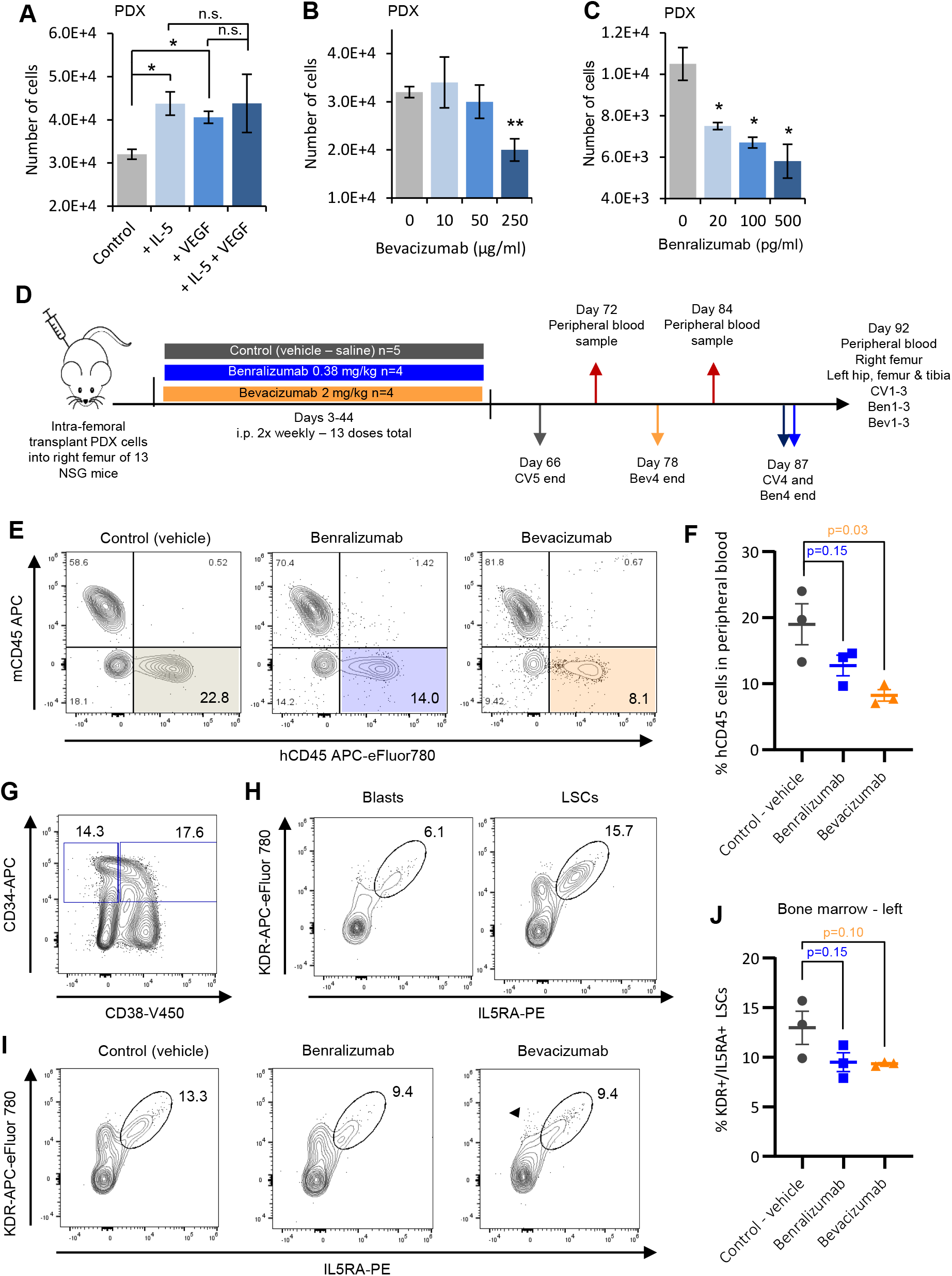
VEGFA and IL5RA inhibitors reduce PDX proliferation. **(A-C)** t(8;21) PDX cells were grown *in vitro* for 6 days with or without IL-5 and/or VEGF(165) **(A),** with 3 doses of bevacizumab **(B)** or with 3 doses of benralizumab (+10ng/ml IL-5) **(C)** and the resulting cells counted. Control/0 bevacizumab sample in A and B is the same as experiments were performed in parallel. Bar height shows the mean of 3 replicates and the error bars indicate SEM. Bar height shows the mean of 3 replicates and error bars indicate SEM. * indicates p <0.05, ** p<0.01, *** p<0.005 using Student’s T-tests. **(D)** Schematic showing how PDX dosing and sampling were conducted *in vivo.* **(E)** Representative contour plots showing the human and mouse CD45 positive cells by flow cytometry in peripheral blood at day 92 post-injection. **(F)** Percentage of human CD45 positive cells in peripheral blood at day 92 post-injection. **(G-H)** Representative contour plots showing the relative populations of hCD45+CD34+CD38+/- cells **(G)** and KDR and IL5RA positivity of hCD45+/CD34+/CD38+ blast cells and hCD45+/CD34+/CD38- LSCs **(H)** in control left bone marrow at day 92 post-injection. **(I)** Representative contour plots showing the KDR and IL5RA positivity of hCD45+/CD34+/CD38- LSCs in treated or control left bone-marrow at day 92 post-injection. **(J)** Percentage of KDR and IL5RA positive hCD45+/CD34+/CD38- LSCs in left bone marrow at day 92 post-injection. **(F & J)** Horizontal and error bars show mean and SEM respectively of the 3 mice in each treatment group, p-values by Student’s t-tests are shown.

These results confirm that t(8;21) patient cells proliferate in response to VEGF and IL-5, preferentially in the LSC compartment. Taken together, these data show that control of LSC growth and self-renewal is mediated by VEGF and IL-5 signalling.

### VEGF and IL-5 signals terminate at the AP-1 family of transcription factors

We then asked how VEGF and IL-5 signalling exert their effects on LSC growth. Both signalling pathways are known to function via MAP-Kinase activation of the AP-1 family of transcription factors to control gene expression, and we and others have shown that AP-1 is a critical regulator of growth and gene expression in t(8;21) AML^7, 18, 20^. To investigate this idea, we generated Kasumi-1^20^ and SKNO-1 cell lines expressing a doxycycline-inducible, flag-tagged, broad range dominant negative FOS (dnFOS) peptide^33^. AP-1 binding to DNA is dependent upon its assembly as a heterodimer of FOS and JUN family proteins coupled via the leucine zipper domains that are adjacent to the basic DNA binding domains. In dnFOS the basic domain is replaced by an acidic domain that binds tightly to the opposing JUN basic domain, thereby blocking binding of all JUN family proteins to DNA. When induced, the peptide was largely localised to the cytoplasm, presumably sequestering JUN proteins before they reach the nucleus (Figure S5A). Combining dnFOS induction with VEGF or IL-5 treatment drastically reduced growth rates compared to the control, negating the stimulation of growth by either factor (Figures 5A-B). The combination of bevacizumab and dnFOS did not show any additive effect (Figures 5A-B). FOS ChIP-seq with Kasumi-1 cells after treatment with bevacizumab showed an almost complete ablation of FOS binding (Figures 5C and S5B). Induction of dnFOS in both t(8;21) cell lines alone significantly reduced the growth rate as compared to an empty vector (EV) control (Figures 5D-E and S5C-D). Furthermore, dnFOS induction significantly reduced colony formation initially but increased comparative re-plating capacity (Figures 5F-G). These results are in concordance with the reduced growth and increased self-renewal seen with bevacizumab and benralizumab treatment and show that blocking AP-1 with dnFOS can be used to simultaneously inhibit both IL-5 and VEGF-stimulated growth. In summary, our data demonstrates that VEGF and IL- 5 signalling controls growth and self-renewal of LSCs upstream of AP-1.

**Figure 5:**
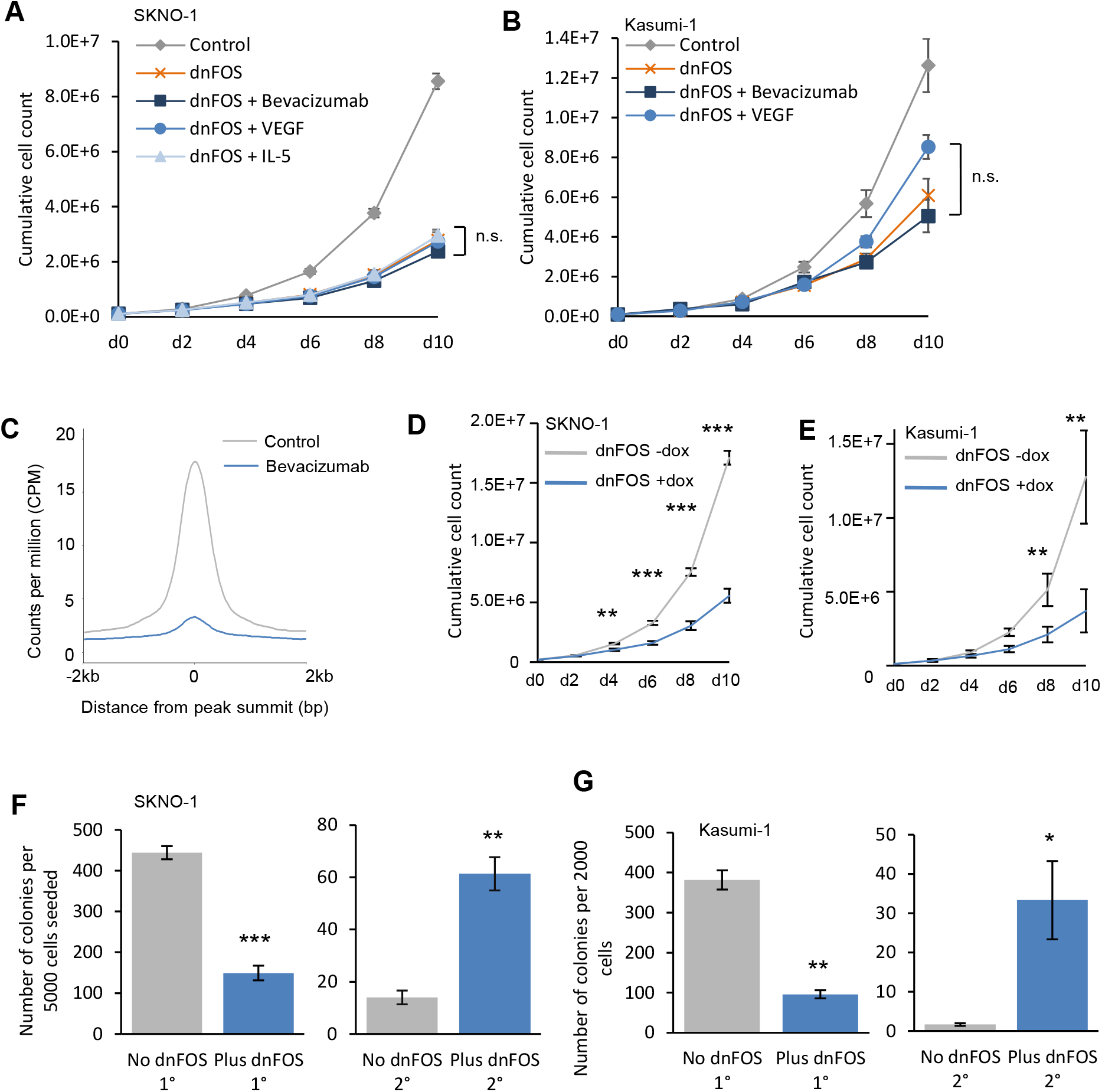
VEGF and IL-5 signals terminate at the AP-1 family of transcription factors (A-B) Growth curves were performed by growing SKNO-1 **(A)** or Kasumi-1 **(B)** cells for 10 days, counting and passaging every 2 days. Cells were grown with induction of dnFOS by doxycycline in conjunction with IL-5 (SKNO-1 only), VEGF-165 or bevacizumab. Each point represents the mean of three experiments, and error bars show SEM. The control curves are the same as in Figure 3 as experiments were performed in parallel and shown again for clarity. No significant differences were found at any time point comparing +dnFOS with any treatment group. **(C)** Histogram showing the average normalised FOS ChIP signal across the union of all peaks, +/-2kb of the summit in Kasumi-1 cells with and without bevacizumab. **(D-E)** Growth curves were performed by growing SKNO-1 **(D)** or Kasumi-1 **(E)** cells for 10 days, counting and passaging every 2 days. Cells were grown with or without dnFOS induced by doxycycline. Each point indicates the mean of three experiments, error bars show SEM. **(F-G)** Colony forming unit assays were performed by plating SKNO-1 **(F)** or Kasumi-1 **(G)** cells in methylcellulose with or without doxycycline to induce dnFOS. The number of colonies were counted after 10 days (left) and cells were replated to form secondary colonies which were again counted after 10 days (right). Bars indicate the mean of three experiments, error bars show SEM. * indicates p <0.05, ** p<0.01, *** p<0.005 using Student’s T-tests.

### AP-1 orchestrates a shift in transcriptional regulation from an LSC to a blast pattern

The data described above suggest that VEGF and IL-5 signalling activate AP-1 to regulate LSC growth. Therefore, we next sought to understand how this circuit feeds into control of gene expression regulating growth. To this end we performed DNaseI-seq, RNA-seq and ChIP-seq experiments for multiple transcription factors (FOS, C/EBP*α*, RUNX1, RUNX1::ETO, PU.1, GATA2) in Kasumi-1 cells with or without dnFOS induction and integrated the data to link AP-1 binding to the wider gene regulatory network. Experiments used a Kasumi-1 cell clone (Figure 6A) expressing high levels of dnFOS in response to doxycycline (Figure S6A). Previously published promoter capture HiC data in the same cell line allowed us to accurately assign distal cis-regulatory elements to their genes^34^.

**Figure 6:**
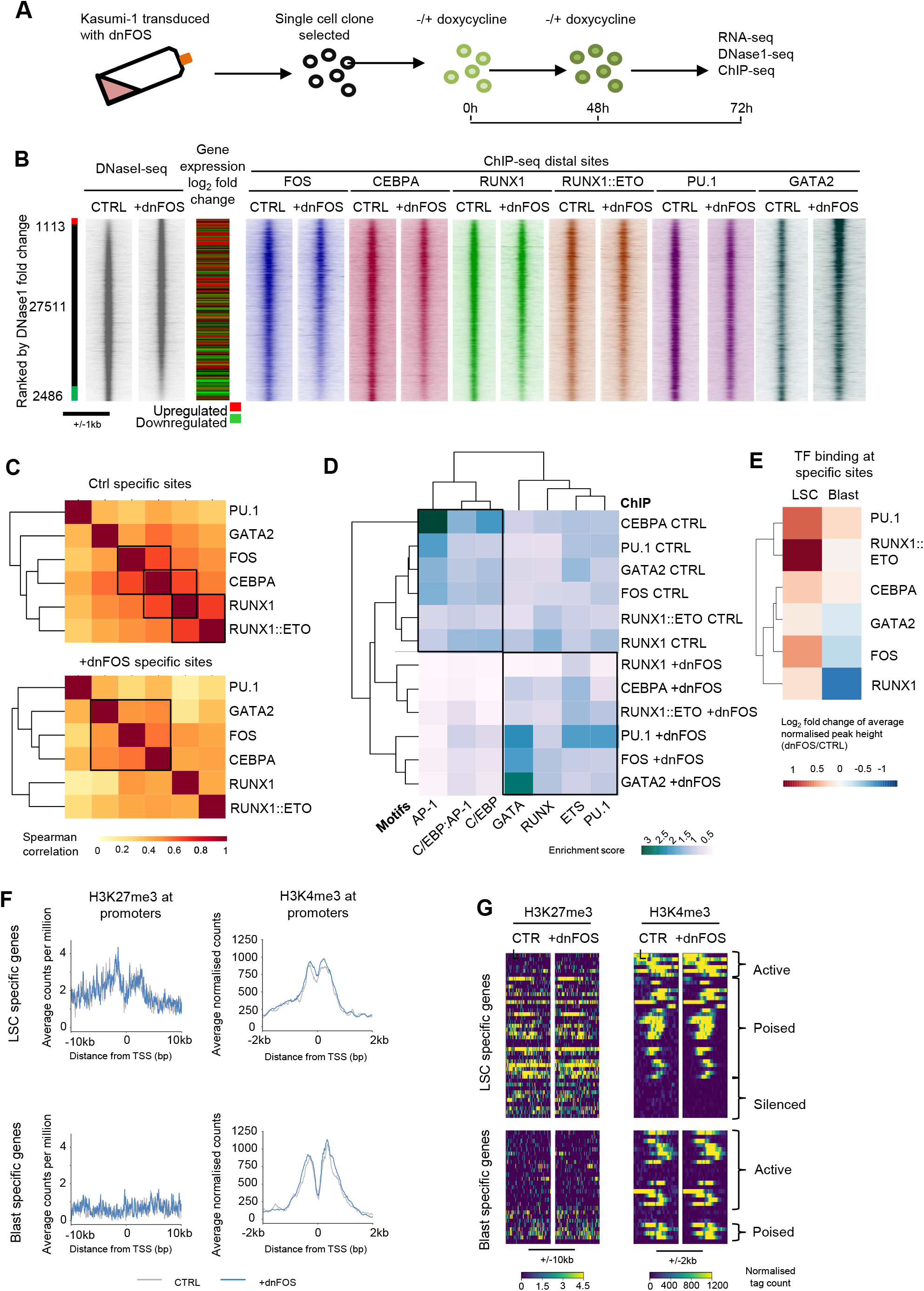
AP-1 orchestrates a shift in transcriptional regulation from an LSC to a blast pattern. **(A)** Schematic showing how dnFOS was induced in primary cells **(B)** DNase1 was performed with and without dnFOS induced by doxycycline in the Kasumi-1 cell line, ranked by fold change of the tag count at distal peaks and represented as density plots (+/1kb of the summit). The red bar indicates dnFOS specific sites and the green bar control specific sites where the normalised tag-count of specific sites is at least two-fold different. ChIP data from FOS, CEBPA, RUNX1, RUNX1::ETO, PU.1 and GATA2 with and without dnFOS were plotted on the same axis across the same window **(C)** Specific sites were calculated for the ChIPs shown in (A) where the normalised tag-count is at least two-fold different in a pairwise comparison of dnFOS against control. The normalised tag count was measured in a peak union generated from control or dnFOS specific sites from all ChIPs and the Spearman correlation calculated which is plotted as a heatmap with hierarchical clustering. **(D)** A motif enrichment score was calculated based on motif frequency in the specific gained (dnFOS) and lost (CTRL) sites calculated in (B) and plotted as a heatmap with hierarchical clustering. **(E)** Heatmap with hierarchical clustering showing the log_2_ fold change between the normalised average peak height of ChIP-seq in Kasumi-1 with dnFOS vs controls at LSC-specific and blast-specific ATAC sites. **(F)** Average profiles were generated from the CPM normalised tag counts of ChIP for H3K27me3 (+/- 10kb from the TSS) and H3K4me3 (+/- 2kb from the TSS), at the promoters of t(8;21) LSC or blast specific genes with or without induction of dnFOS in Kasumi-1 cells. **(G)** Density plots showing the signal at each of the sites used in **(F)**, with active, silenced and poised genes indicated.

The comparison of LSC and blast open chromatin regions had shown that LSC-specific accessible chromatin sites were enriched in GATA motifs (Fig S1D). We noted that DNaseI-seq in Kasumi-1 cells expressing dnFOS showed gain of chromatin accessibility associated with increased binding of GATA2 mostly at distal chromatin sites (Figures 6B and S6B). Lost chromatin accessibility was associated with loss of binding of FOS, RUNX1, RUNX1::ETO, C/EBPα and PU.1. To ask whether these factors are binding in combination and therefore were jointly lost directly due to blockade of FOS binding, we examined the ChIP signal across a union of gained and lost binding sites for all six factors and performed a correlation analysis of the tag counts (Figure 6C). This analysis examines whether the gained and lost peaks are the same for each factor, and indicated highly correlated binding patterns of RUNX1, RUNX1::ETO, FOS and C/EBPα at lost sites (Ctrl) and correlation of GATA2, C/EBPα and FOS binding at gained sites (Figure 6C). Furthermore, an analysis of the motif spacing in the lost sites in each ChIP showed similar proximity of the RUNX1, RUNX1::ETO and C/EBP consensus sequence to the AP-1 motif (Figure S6C). Motif enrichment analysis confirmed that the gained sites (dnFOS) were particularly enriched for GATA, but not AP-1 motifs, whilst the lost sites (CTRL) shared AP-1, C/EBP and composite C/EBP:AP-1 motifs (Figure 6D). C/EBP and AP-1 family members can heterodimerise^35^ and *CEBPA* is repressed in t(8;21) AML. We therefore queried whether this result indicated that AP-1:C/EBP heterodimers were being disrupted, and if this was a facet of the importance of AP-1 in t(8;21) AML. We therefore expressed a dnCEBP peptide in an inducible fashion as well^12, 33^. C/EBP is required to maintain the viability of Kasumi-1 cells and whilst dnCEBP expression led to loss of open chromatin containing AP-1, C/EBP, CEBP:AP-1 composite and RUNX1 binding motifs, it did not lead to gain of sites associated with GATA binding but instead gained AP-1 and RUNX1 binding sites (Figure S6D)^12^. Directly comparing the sites which were lost and gained with dnFOS and dnCEBP revealed that sites which were gained with dnFOS also gained accessibility with dnCEBP, and sites lost with dnCEBP also lost accessibility with dnFOS (Figure S6D) but the overlap was incomplete (Figure S6E) underpinning the discrepancy in motif patterns. Therefore, dnFOS and dnCEBP do not impact upon the same aspects of gene regulation, and whilst loss of AP-1 binding is associated with loss of C/EBPα binding, loss of AP-1 activity specifically contributes to a gain of GATA2 binding.

After induction of dnFOS expression, 226 genes were significantly up-regulated and 60 were down- regulated by at least 2-fold (Figure S6F, Table S3). The comparison of binding alterations as measured by ChIP and gene expression changes showed that loss of FOS, RUNX1 and RUNX1::ETO binding led to both up and down-regulation of genes (Figure S6G). However, the acquisition of GATA2 binding was predominantly observed at elements associated with up-regulated genes (white shows lack of binding at an element). *GATA2* expression was also up-regulated, with increased binding to its enhancers thus setting up an auto-regulatory loop (Figures S6H and S6I). GATA2 is a key regulator of stem cell maintenance^23^ and we and others have shown that GATA2 is expressed in LSCs (Figure 1)^36^. To examine whether the increase in GATA2 binding after AP-1 inhibition would lead to a reactivation of LSC-specific genes we first examined transcription factor binding at LSC or blast cell specific genes as defined in Figure 1 with and without dnFOS induction in Kasumi-1 cells (Figure 6E). This analysis showed a relative reduction in binding of FOS, RUNX1 and GATA2 at blast associated sites, whilst GATA2, RUNX1::ETO, PU.1 and FOS binding was increased at LSC-specific sites in the absence of AP-1 activity (Figure 6E). We then investigated histone modifications associated with promoter activation and silencing at the LSC and blast specific genes. This analysis showed that in the Kasumi-1 cell line LSC-specific gene promoters were bound by H3K27me3 (Figure 6F). Moreover, LSC promoters were also marked with H3K4me3 indicating that they are in a bivalent or poised chromatin conformation^37^. Together these data show that inhibiting AP-1 leads to reactivation of poised LSC genes in blast cells, and a silencing of blast genes through a shift in FOS and PU.1 to GATA2 sites and loss of RUNX1 particularly from AP-1 and C/EBP sites.

### AP-1 is required for maintenance of the blast program

To confirm the notion that inhibition of AP-1 restores an LSC signature in primary t(8;21) AML cells, we transduced dnFOS or an empty vector control into PDX cells and into healthy CD34+ PBSCs. We sorted the GFP expressing cells following transduction and dox induction resulting in a population expressing either GFP alone or dnFOS (Figures 7A and S7A). In PDX cells 160 genes were up- regulated and 129 genes down-regulated by at least 2-fold whilst in the healthy cells fewer genes were de-regulated (Figures 7B and 7C, Table S4). In concordance with this result, healthy cells did not show a phenotypic response to dnFOS in colony forming assays (Figure S7B). To ask whether dnFOS induction impacted the LSC and blast gene expression programs, we performed GSEA based on the joint t(8;21) LSC and blast genes (Figure 1C), as well as the LSC and blast specific genes from each patient respectively. In all cases, the genes up-regulated in the PDX cells expressing dnFOS were strongly enriched for LSC genes, and the down-regulated genes for the blast signature, which was not the case for healthy PBSCs (Figures 7D-F and S7C-F).

**Figure 7:**
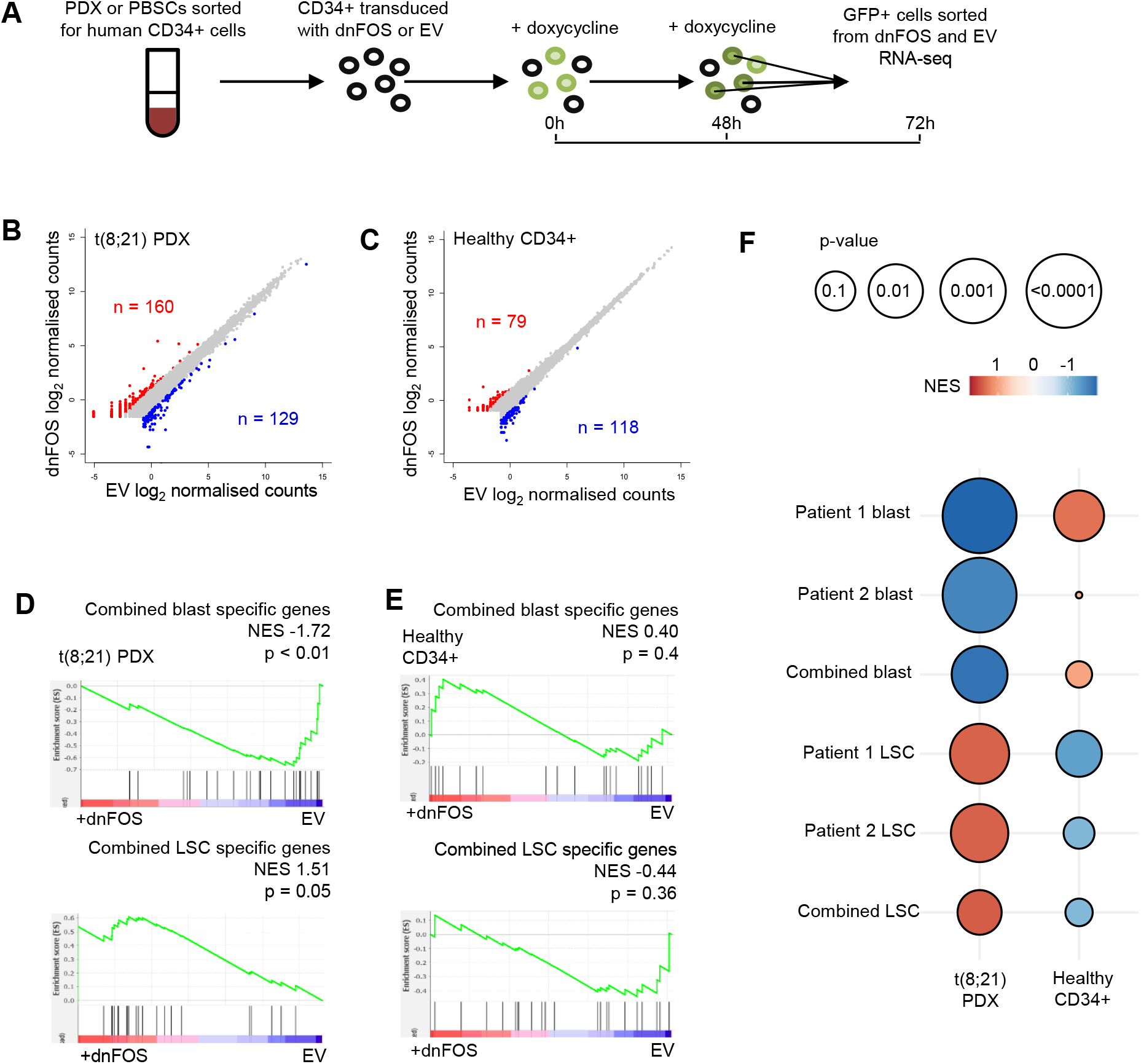
AP-1 is required for maintenance of the blast program. **(A)** Schematic showing how dnFOS was induced in primary cells **(B-C)** RNA-seq was performed in PDX cells and healthy CD34+ PBSCs following induction of dnFOS or the EV control, gene expression is shown as a scatter plot of the log_2_ counts, with the genes up-regulated by dnFOS highlighted red and the down-regulated genes highlighted in blue. **(D-E)** GSEA was used to compare blast and LSC specific genes identified in 1C with the ranked fold change gene expression from the PDX **(D)** and healthy CD34+ cells **(E)**, comparing dnFOS to EV. NES shows the normalised enrichment score from the GSEA and the adjusted p-value. **(F)** Bubble plot showing the results of GSEA in D and E, as well as with each individual patients’ LSC and blast specific genes. The colour scale indicates the normalised enrichment score, whilst the size of the bubble indicates the adjust p-value, where a larger bubble is more significant.

### The signalling response of t(8;21) cells operates within a RUNX1::ETO dependent regulatory circuit

RUNX1::ETO is required for the maintenance of the leukemic state in t(8;21) cells as its depletion activates a C/EBPα-dependent myeloid differentiation program^14, 15, 34, 38^. The above data show that AP-1 is also required to support growth of t(8;21) cells but AP-1 family member gene expression is a feature of most subtypes of AML (Figure S8A). Of the six AP-1 family genes most highly expressed in t(8;21) AML we found that all were expressed at significantly higher levels in LSCs compared to blasts in patient 1, and all except FOSB showed significantly higher expression in LSCs and/or the LSC transition population in patient 2 (Figure S8B). Furthermore, the expression of RUNX1::ETO leads to the activation of *JUN* expression^17, 39, 40^. We therefore asked how the VEGF and IL-5 signalling pathways, together with AP-1 family members, are regulated with respect to the driver oncoprotein RUNX1::ETO by investigating gene expression and the genomic landscape with and without RUNX1::ETO depletion in a Kasumi-1 cell line carrying an inducible shRNA targeting RUNX1::ETO. The knockdown experiments showed that *JUNB*, *JUN* and *JUND* were up-regulated in the presence of RUNX1::ETO, whilst expression of their partners *FOS* and *FOSB* was not (Figure 8A). *VEGFA* was also down-regulated with RUNX1::ETO knockdown suggesting that both the specific signalling molecules and the downstream regulators are all influenced by the presence of the driver oncoprotein (Figure 8A). FOS shows a large overlap in binding sites with JUN and JUND^34^ in wild-type Kasumi-1 cells as shown by ChIP-Seq (Figure S8C). However, JUN and FOS proteins behaved differently with respect to RUNX1::ETO depletion, as exemplified by FOS and JUND. JUND binding and expression were decreased after knockdown of RUNX1::ETO (Figures 8A and 8B). In contrast, FOS was lost from distal cis-regulatory elements containing AP-1 motifs and re-distributed to promoters with accessible chromatin and bound PolII (Figures 8B, S8D and S8E). Most FOS and RUNX1::ETO binding sites, whilst responsive to oncoprotein depletion, do not overlap. However, many FOS sites do overlap with RUNX1 binding (Figure S8F). Together these data show that both AP-1 expression and localisation are orchestrated by RUNX1::ETO and further regulated by VEGF and IL-5 signalling. We next asked how RUNX1::ETO, AP-1 and the response to VEGF and IL-5 signalling are integrated into control of the chromatin landscape underpinning leukemogenesis. We found that the signalling responsive histone modification H3K9acS10P^41^ was dramatically and globally reduced following RUNX1::ETO knockdown despite an increase in total H3K9ac (Figure S8G)^14^. Although not directly associated with the altered FOS binding the loss of this histone modification implies signalling to chromatin indeed relies upon RUNX1::ETO.

**Figure 8:**
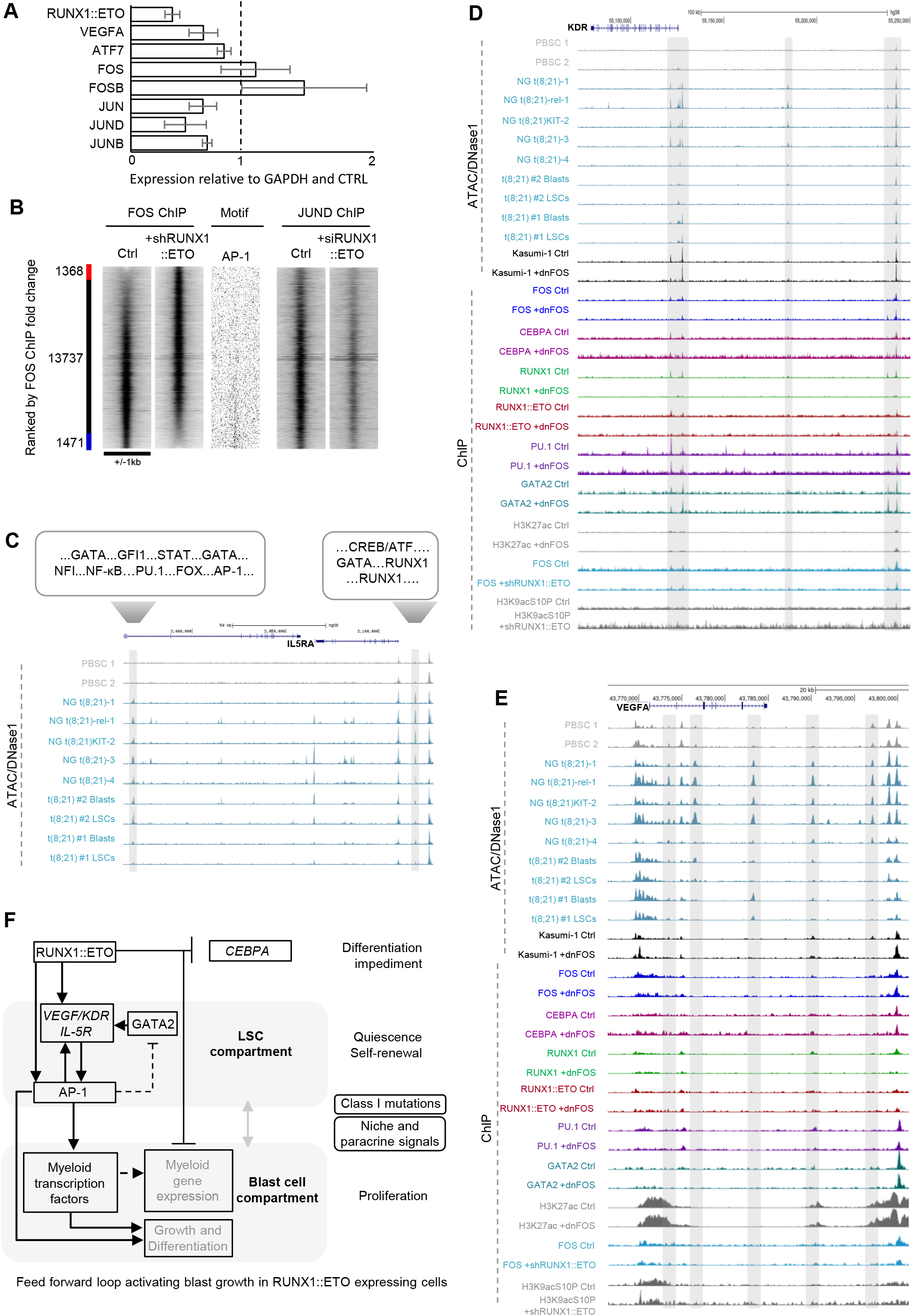
The signalling response of t(8;21) cells operates within a RUNX1::ETO dependent regulatory circuit. **(A)** qRT-PCR showing the relative change in expression of *RUNX1::ETO, VEGFA* and the most highly expressed AP-1 members after shRUNX1::ETO knockdown. Bars indicate the average of 4 replicates, error bars show SEM, the vertical dashed line indicates no change in expression. **(B)** ChIP for FOS was performed with and without shRUNX1::ETO induced by doxycycline in the Kasumi-1 cell line, ranked by fold change of the tag count at all peaks and represented as density plots (+/1kb of the summit). The red bar indicates shRUNX1::ETO specific sites and the blue bar control specific sites where the normalised tag-count of specific sites is at least two-fold different. ChIP for JUND with siMM (Ctrl) or siRUNX1::ETO^34^ and AP-1 motif frequency is plotted along the same axis across the same window. **(C-E)** UCSC genome browser screenshots showing ATAC/DNaseI in healthy CD34+ PBSCs and t(8;21) AML patients^7^ at the *IL5RA* locus, with the transcription factor binding motifs in the t(8;21) specific peaks indicated **(C)**, and additionally showing DNaseI and ChIP in Kasumi-1 +/- dnFOS, and +/- shRUNX1::ETO at the *KDR (D)* and *VEGFA (E)* loci with the t(8;21) specific peaks indicated. *(f)* Model showing how AP-1 activated by signalling activates blast cell growth in t(8;21) AML.

AP-1 gene expression and its binding to DNA are normally only detected at a substantial level in the presence of active signalling^42^. To investigate whether the expression of *VEGFA* and *IL5RA* is signalling or RUNX1::ETO responsive and thus form a regulatory circuitry, we examined their cis-regulatory regions. We assembled DNaseI-seq data from t(8;21) AML patients together with the above-described DNaseI-seq and ChIP-seq data for myeloid transcription factors from Kasumi-1 cells in the presence or absence of dnFOS. Two results were noteworthy: (i) All three genes showed a DHS at their promoters in healthy PSBCs (Figures 8C-E) and in purified HSCs^22^ suggesting that their promoters were still poised for expression; (ii) the *VEGFA* and *KDR* promoters were bound by FOS whose binding was responsive to dnFOS and RUNX1::ETO depletion, linking gene expression control directly to factor binding.

At the *IL5RA* locus, two specific DNaseI peaks were detected in t(8;21) AML patients (Figure 8C, indicated by a grey bar) but not in healthy PBSCs. No ChIP-seq signal was detected here, due to the likely loss of this region of chromosome in Kasumi-1 cells. A motif search in the DNaseI hypersensitive sites (DHSs) from primary cells revealed GATA, PU.1, FOX, AP1, CREB/ATF and RUNX binding motifs (Figures 8C and S8H). Regulation of *VEGFA* and *KDR* was more complex, with multiple peaks and broad regulatory regions (Figures 8D and 8E). None of the DHSs were exclusive to LSCs, suggesting that signalling-responsive transcription factor binding activity controls specificity of expression. After inhibition of AP-1 by dnFOS we indeed observed loss of chromatin accessibility and FOS binding at these DHSs, and at some peaks loss of RUNX1 binding as well. After shRUNX1::ETO induction, both FOS binding and the H3K9acS10P were largely unchanged at these sites, despite VEGFA expression going down after RUNX1::ETO knockdown (Figure 8A). Taken together, our data show that *VEGFA,* KDR and *IL5RA* are regulated by a complex interplay of activating and repressing transcription factors operating within the context of a primed and signalling responsive chromatin landscape.

## Discussion

In this study, we have shown that t(8;21) LSCs specifically utilise VEGF and IL-5 signalling to promote growth. VEGF and IL-5 are aberrantly expressed in t(8;21) LSCs as part of a regulatory circuit involving the driver oncogene RUNX1::ETO and the AP-1 complex as a mediator of signalling. This interplay forms a feed-forward loop with RUNX1::ETO at the apex (Figure 8F). RUNX1::ETO blocks differentiation by down-regulating *CEBPA*^43^ and disrupting PU.1 and RUNX1 driven control of myelopoiesis^14, 44^. Simultaneously, RUNX1::ETO, when expressed on its own, blocks the cell cycle^17^ which is overcome via AP-1 dependent gene regulation^20^. Moreover, AP-1 is also required for myeloid differentiation as its inhibition up-regulates *GATA2* expression and shifts cells to a more immature state. *JUN* is up-regulated in an indirect but RUNX1::ETO dependent fashion^40^ whereas *FOS* expression is controlled by outside signals as well^45^ but expression itself does not respond to RUNX1::ETO. In summary, the acquisition of signalling mutations and/or signals from the niche cause the up-regulation and post-translational activation of the AP-1 complex. AP-1 then orchestrates changes in the transcriptional program through altering CEBPA, PU.1, RUNX1 and GATA2 binding patterns, leading to a reversible silencing of LSC genes and the activation of blast genes, preserving self-renewal capacity whilst allowing cell expansion.

Like healthy HSCs, LSCs mostly remain quiescent, allowing them to evade chemotherapy, despite the presence of constitutively active mutant cytokine receptors which drive proliferation of blasts. The expression of RAS mutations and likely also mutant growth factor receptors activating the RAS pathway is detrimental to HSC maintenance even in the presence of RUNX1::ETO^46^. Thus, LSCs carrying such mutations are likely not to arise from HSCs, but instead co-opt the chromatin profile and gene expression patterns of such cells^5, 47^, including expression of GATA2 and JUNB which control the regulation of cell cycle genes^48–50^. The fact that LSCs develop an HSC-like regulatory phenotype may be part of a chemotherapy response^51^ or occur because of the absence of specific growth signals at their location, as mimicked by the dnFOS experiments shown here. We have shown here that LSC-specific genes are marked with bivalent chromatin meaning they are not irreversibly silenced^52–54^, and can be re-activated when signals to LSCs are blocked, forming a feedback loop that prepares cells to respond to extrinsic signals.

*VEGFA, KDR* and *IL5RA* are not normally expressed in myeloid or stem cells but show a primed chromatin structure in HSCs with the promoters being still hypersensitive and ready to be expressed^22^. Each of these genes is a target for AP-1 mediated signalling transduction in established AML cells, but AP-1 is also involved in co-opting *VEGFA* into supporting the growth of non-myeloid leukemic cells^55^. *VEGFA*, *KDR* and *IL5RA* are also GATA2 targets, whereby GATA2 further co-operates with AP-1^56^ and is specifically expressed in LSCs which are poised to cycle^57^. An important result from our study is therefore that the exact signalling pathways employed by LSCs are highly subtype specific, relying on the specific interplay of the driver mutation with the stem cell program. Using published scRNA-seq data we can confirm this notion, with a cluster of cells co-expressing *IL5RA*, *VEGFA* and *GATA2* detected in the t(8;21) AML sample^58^. During embryonic development and thereafter VEGFA and KDR which are part of the endothelial gene expression program are repressed by RUNX1^59^, RUNX1::ETO disrupts the action of wild-type RUNX1 on VEGFA/KDR^60^ and endothelial gene expression remains elevated^61, 62^. In biCEBPA mutant AML, RUNX1 expression is down- regulated^12^ as well and as a result *VEGFA, KDR* and *IL5RA* are still expressed but at a lower level than in t(8;21). However, note that the shared IL-3/IL-5/GM-CSF receptor CSF2RB is specifically enriched in GATA2-high biCEBPA-mutant LSCs^12^. Moreover, inspection of LSC and blast cell single cell data demonstrated that also this type of AML activates a specific, but different growth factor / receptor pair suggesting that pathway hijacking is used by more than one AML sub-type^12^.

Activation of mis-expressed signalling pathways in LSCs leads to the re-generation of full-scale leukemia with the signals coming from the environment in which they reside. IL-5 is normally produced by eosinophils, mast cells and stromal cells, whilst VEGF-signalling is coming from the vascular niche as well as from AML cells themselves. VEGF also contributes to engineering of the niche by leukemic cells to better support their growth^63–65^. In this scenario, relapse is inevitable as LSCs are ready and waiting for the signals which will eventually arrive. It has been shown that LSCs undergo a transient amplification after chemotherapy^51^. Therapy therefore needs to simultaneously target rapidly growing blast cells and block signalling to prevent re-entry of LSCs into the cell cycle. In t(8;21) AML this may be achieved by repurposing the FDA approved monoclonal antibodies bevacizumab and/or benralizumab. Inhibition of VEGF by bevacizumab has been previously trialled in AML to block remodelling of the niche but only 2 core-binding factor AML patients of unknown genotype were included^66^ and the results overall were therefore inconclusive. In summary, our work highlights the importance of studying the fine details of AML sub-type specific gene regulatory networks impacting on specific mechanisms of growth control to find the right therapeutic targets to prevent relapse.

## Acknowledgements

This work was funded by grants to C.B. and P.N.C. from Blood Cancer UK (20006), a grant from the Medical Research Council (MR/S021469/1) to C.B., P.N.C and O.H, a Cancer Research UK programme grant (C27943/A23389) and a KIKA programme grant (329) to OH, a Leukemia UK John Goldman Fellowship (2021/JGF/001) to D.J.L.C and a Cancer Research UK studentship to A.A. Benralizumab was obtained from Astra Zeneca in the context of the Open Innovation scheme. The authors would like to acknowledge Celina Whalley of Genomics Birmingham and Guillaume Desanti of the University of Birmingham Flow Cytometry facility for support of next-generation sequencing and cell sorting experiments. We would like to thank the West Midlands Regional Genetics Laboratory for supplying mutation data linked to AML patient samples. We would like to thank Hesta McNeill and Samantha Jepson Gosling of Newcastle University for their technical assistance.

## Author contributions

S.G.K and C.B. conceived and directed the work and wrote the paper, S.G.K., S.P., H.J.B., P.S.C, A.P., A.W., L.A. A.A., D.J.L.C, N.K., A.K-H. performed experiments, M.R. and S.P. provided patient cells, S.G.K., P.K and S.A.A. analysed data, P.N.C. helped writing the paper, H.J.B. and O.H. directed the mouse work and helped writing the paper.

## Conflict of Interest

The authors declare no conflict of interest.

## Methods

### Key resources table

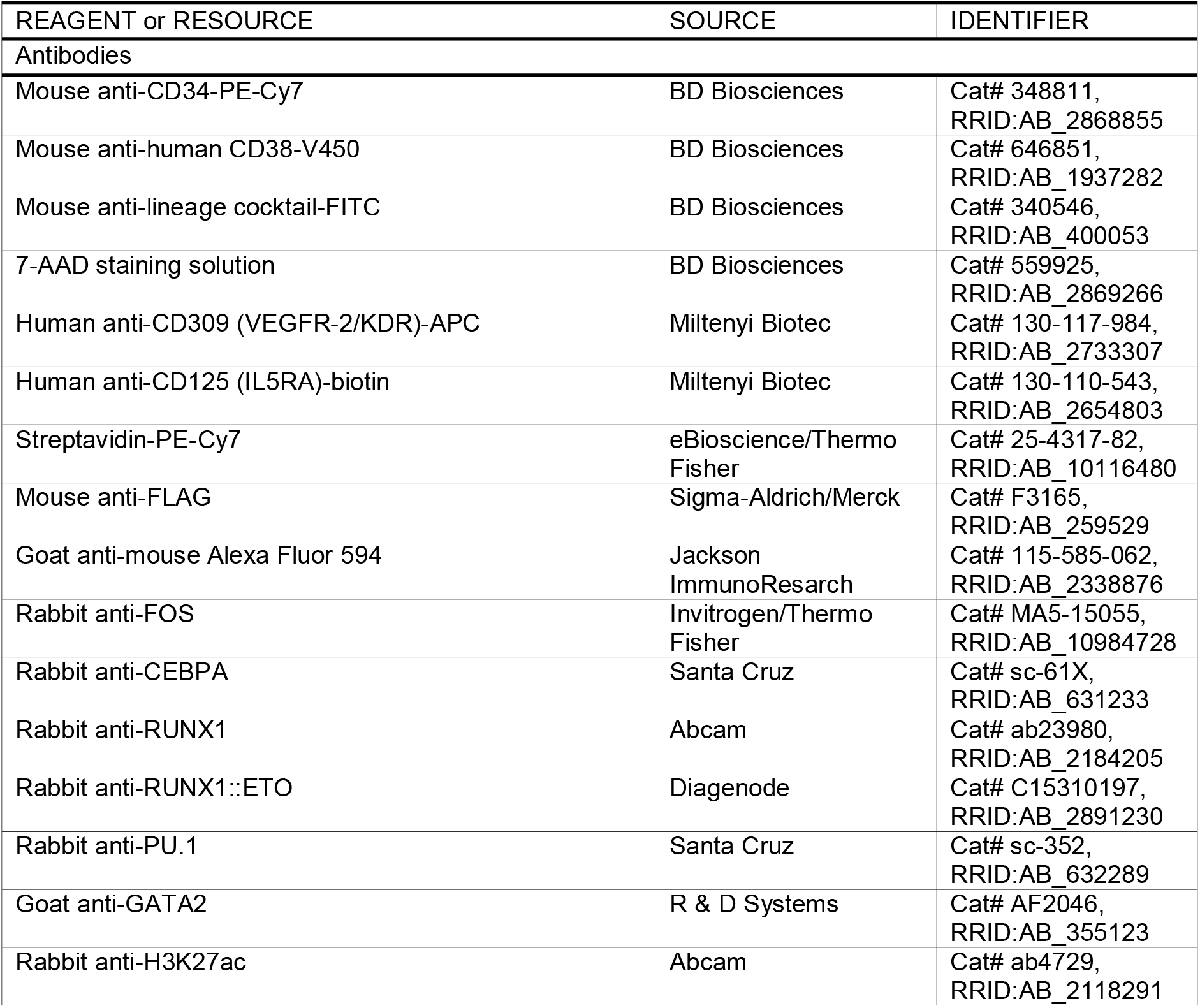
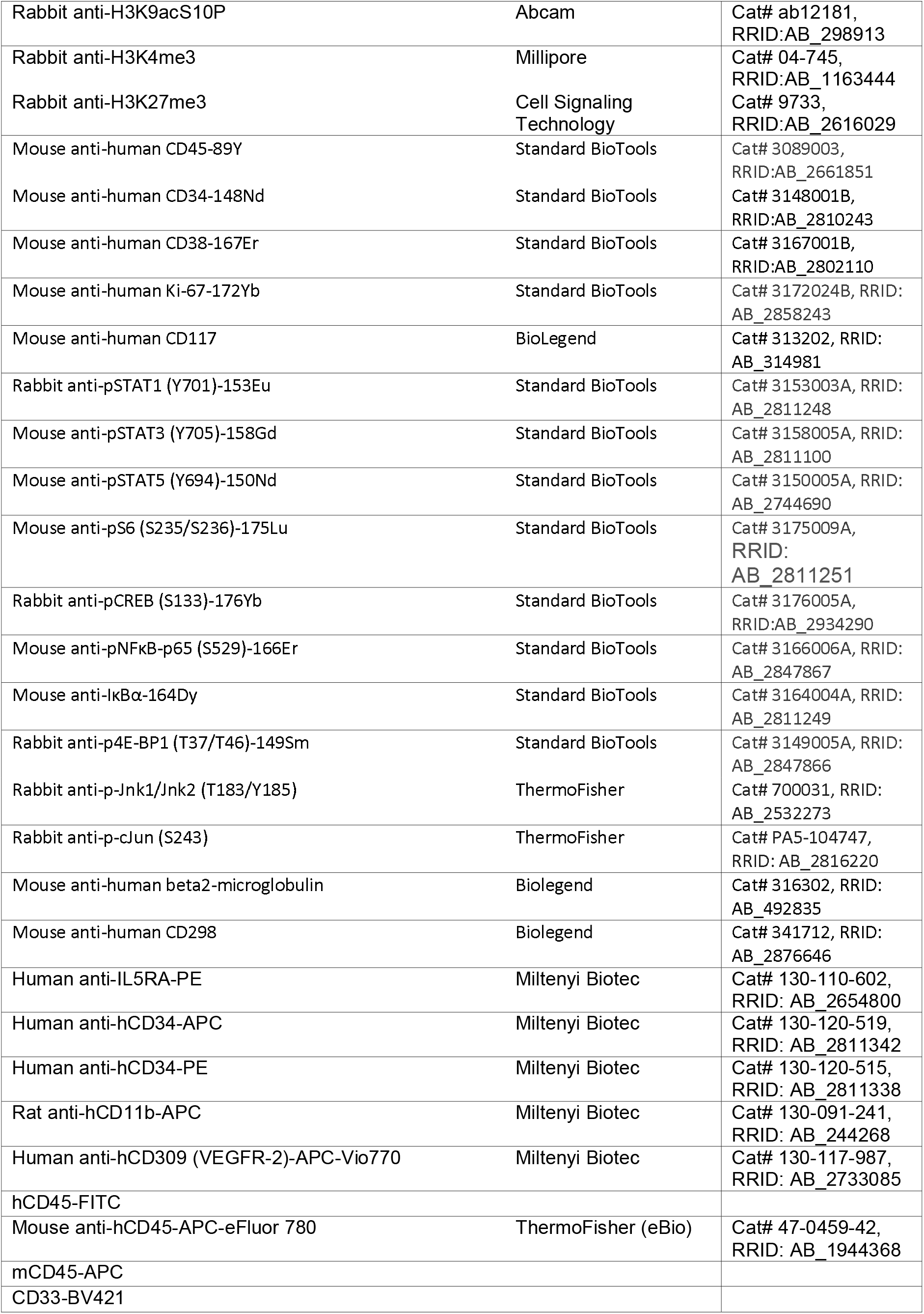
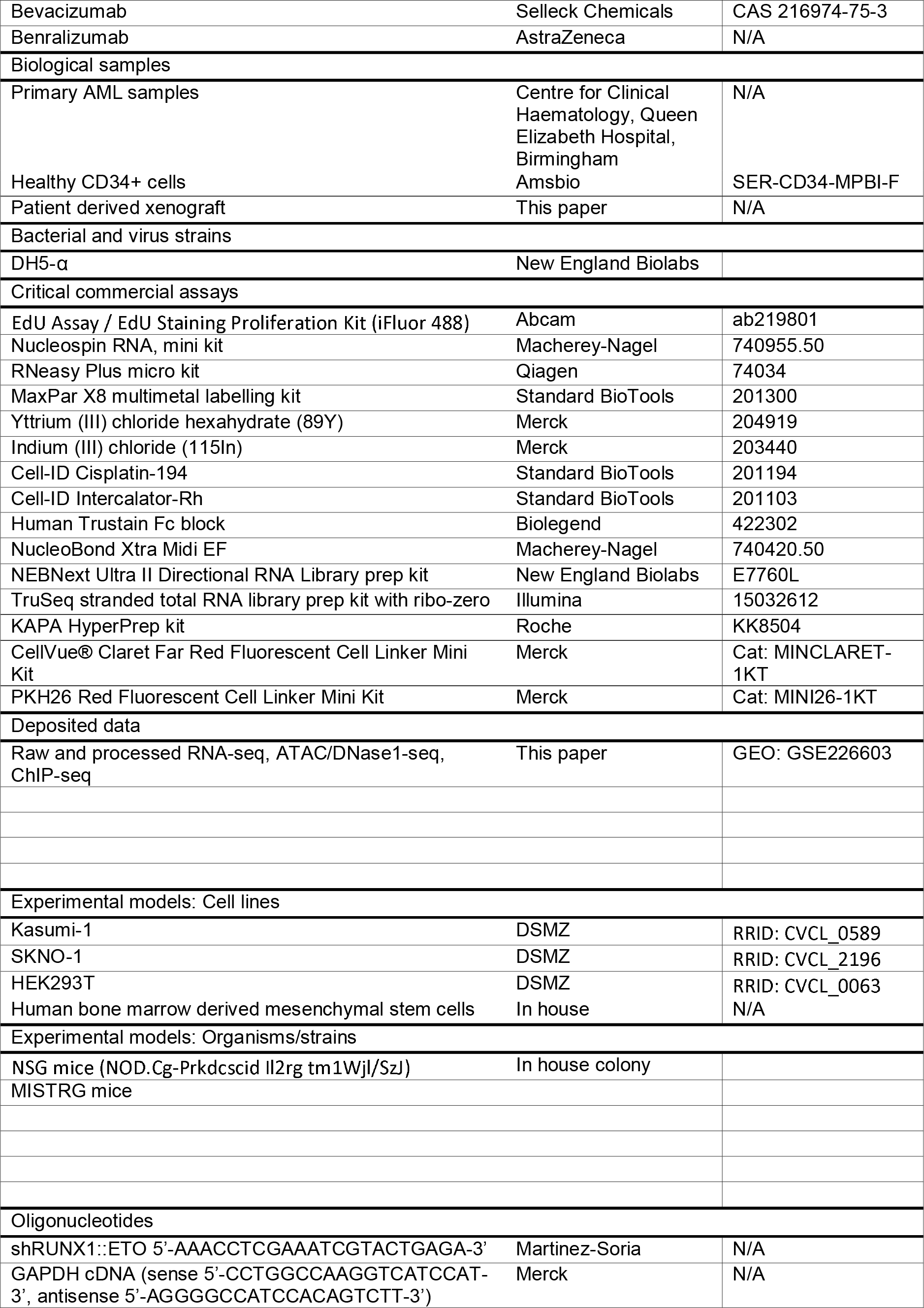
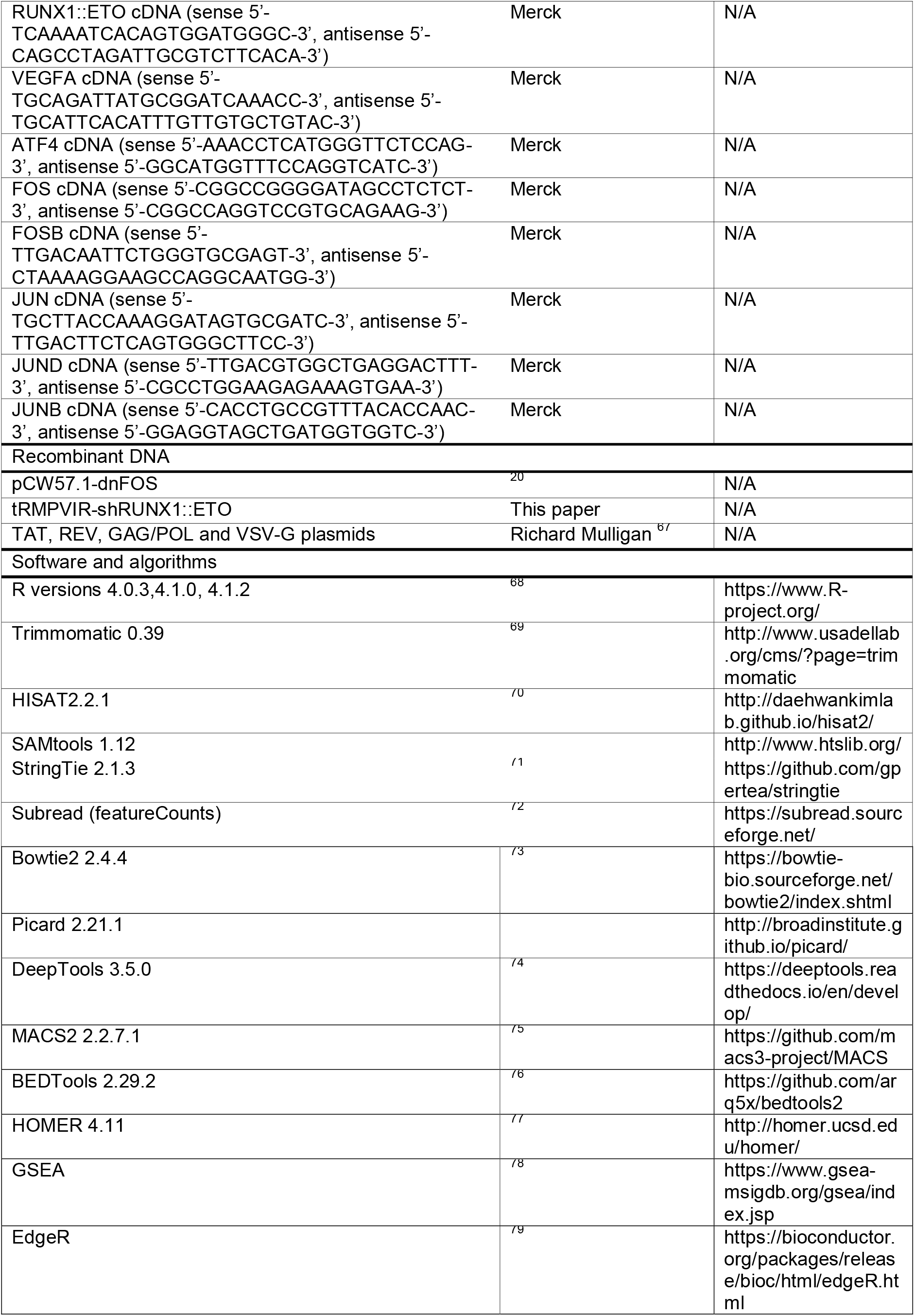
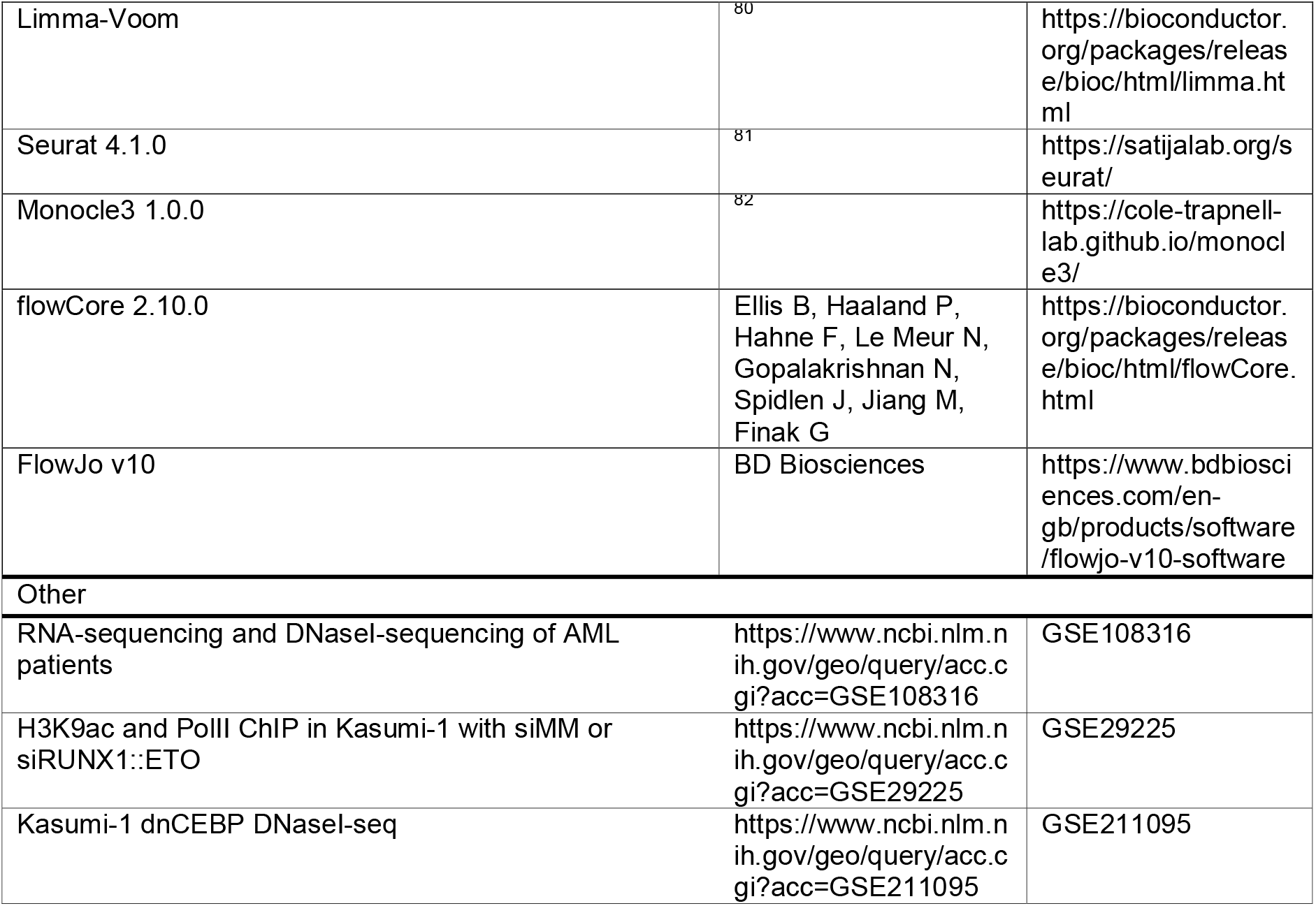

### Resource availability

#### Lead contact

Further information and requests for resources should be directed to the lead contact, Constanze Bonifer.

#### Materials availability

Plasmids generated in this study are available by request from the lead contact.

#### Data and code availability

RNA-seq, ATAC/DNase-seq, ChIP-seq and scRNA-seq have been deposited at Gene Expression Omnibus publicly available as of the date of publication.

Accession numbers are listed in the key resources table.

The paper does not report original code.

Any additional information required to reanalyze the data reported in this paper is available from the lead contact upon request.

### Experimental model and subject details

#### Cell lines

Kasumi-1 cells (RRID: CVCL_0589; male) and SKNO-1 cells (RRID: CVCL_2196; male) were routinely maintained in RPMI1640 medium supplemented with 10% or 20% FBS respectively, 2mM L-Glutamine and 1% Penicillin/Streptomycin. SKNO-1 cells were additionally supplemented with 10ng/ml GM-CSF. HEK293T cells (RRID: CVCL_0063; female) were maintained in DMEM with 10% FBS, 2mM L-Glutamine and 1% Pencillin/Streptomycin. All cells were incubated at 37°C in a humidified 5% CO_2_ incubator.

#### Primary cultures

Patient bone marrow biopsies were obtained, and the AML cells purified using lymphoprep followed by CD34 MACS bead enrichment. Patient mutation details are in Table 1. Primary cells and patient-derived xenograft cells (patient 4 only) were cultured on human mesenchymal stem cells, in SFEMII (StemCell Technologies) supplemented with 1% Pencillin/Streptomycin, 1 µM UM729 (Stemcell Technologies), 750nM SR1 (Stemcell Technologies), 150ng/ml SCF, 100ng/ml TPO, 10ng/ml FLT3, 10ng/ml IL3, 10ng/ml GM-CSF (all cytokines from Peprotech). Where primary cells were frozen prior to use, they were allowed to recover for a week before performing phenotypic assays but sorted directly from defrost for gene expression analysis. Healthy CD34+ cells (Amsbio) were cultured in SFEMII with StemSpan CD34 Expansion Supplement (Stem Cell Technologies) and 500nM UM729 for 1 week, then moved into the t(8;21) media for 24 hours prior to setting up assays.

#### In vivo Experiments

All mouse studies were carried out in accordance with UK Animals (Scientific Procedures) Act, 1986 under project licence P74687DB5 following approval from Newcastle University animal ethical review body (AWERB). Mice were housed in specific pathogen free conditions in individually ventilated cages with sterile bedding, water and diet (Irradiated RM3 breeding diet, SDS). All procedures were performed aseptically in a laminar flow hood. NSG mice (NOD.Cg-Prkdcscid Il2rg tm1Wjl/SzJ) aged between 12 and 16 weeks, both sexes, from an in-house colony were transplanted intra-femorally under isoflurane anaesthetic and 5mg/kg subcutaneous NSAID analgesia (Carprofen). Newborn MISTRG mice were injected intra-hepatically according to Ellegast et al.^83^ Mice were checked daily, weighed and examined at least once weekly to ensure good health. Endpoints for humane killing were pale extremities, hunched posture, 20% weight loss compared to highest previous weight or 10% weight loss for 3 consecutive days and tumours of 1.5cm diameter.

### Method details

#### Plasmid generation

Generation of the dnFOS plasmid was previously described^20^ - dnFOS was amplified from cDNA provided by Charles Vinson^33^ with SalI and NotI restriction site overhangs. Using these restriction sites, the fragment was ligated into pENTR2B (Addgene) and then recombined into pCW57.1 (Addgene). The empty vector was pCW57.1 alone. The shRUNX1::ETO plasmid was generated with XhoI and EcoRI restriction site overhangs. Using these restriction sites, the fragment was ligated into tRMPVIR (Addgene) plasmid. Plasmids were selected and propagated in DH5α competent cells prior to maxiprep using NucleoBond Xtra Midi EF kit and then lentiviral production.

#### Lentivirus production and cell transduction

Lentivirus was produced in HEK293T cells using calcium phosphate co-precipitation of the target plasmid and packaging vectors TAT, REV, GAG/POL and VSV-G at a mass ratio of 24 μg : 1.2 μg : 1.2 μg : 1.2 μg : 2.4 μg per 150 mm diameter plate of cells. Viral supernatant was harvested after 24, 36, 48 and 60 hours then concentrated by ultracentrifugation at 25,000xG for 1 hour 45 minutes at 4°C. Concentrated virus was then transduced into cell lines or primary cells with 8 µg/ml polybrene via spinoculation at 1500xG for 45 minutes. Media was refreshed after 12 hours. To generate clones, cell lines underwent puromycin selection (1 µg/ml) for 5 days and were then sorted for single cells by FACS.

#### Growth curves

For growth curves, cell lines were counted using trypan blue and passaged every 2 days, seeding cells at the original concentration. Cells were grown with 10 ng/ml IL-5 (Peprotech), 50 ng/ml VEGF- 165 (Peprotech), 10 µg/ml Bevacizumab (Selleckchem) and/or 500 pg/ml Benralizumab (AstraZeneca). Where appropriate, doxycycline induction of transduced cells was at 2 µg/ml. Growth curves were not performed in the PDX, instead the cells were just counted at day 6 after seeding.

#### Colony forming assays

For colony assays, cells were grown for 24 hours with the treatment to be tested, then seeded into H4100 MethoCult (Stem Cell) with RPMI1640 and 10% FBS, and the treatment to be tested including doxycycline as appropriate. Patient-derived cells were seeded into MethoCult Express (Stem Cell) Kasumi-1 were seeded at 2000 cells per dish, SKNO-1 were seeded at 5000 cells per dish and patient cells were seeded at 1000 cells per dish. Colony assays were counted after 10 days, except for patient-derived colonies which were assessed after 20 days.

#### Flow cytometry/FACS

Flow cytometry was carried out on a Cyan ADP (Beckman Coulter) using antibodies against CD309- APC (KDR) and CD125-biotin (IL5RA) followed by streptavidin-PE-Cy7 for cell lines, and on an Attune NxT (Thermo Fisher) using antibodies against 1: hCD45-FITC, CD34-APC, CD38-V450, VEGFR-APC- vio770 and IL5RA-PE, 2: hCD45 APC-eFluor780, CD34-PE and mCD45-APC, or 3: CD33-BV421, CD11b- APC, CD34-PE and hCD45-APC-eFluor780 for PDX cells. Cells were resuspended in 100 µl MACS buffer (PBS + 2mM EDTA + 0.5% BSA) and all antibodies were added at 1:100, with staining for 30 mins at 4°C. Compensation was set up using cells and/or compensation beads. Analysis was carried out on FlowJo v10.

FACS was carried out using a FACS Aria (BD). LSCs and blasts were identified and sorted using 7-AAD and lineage cocktail-FITC to select lineage-negative viable cells, followed by CD34-PE-Cy7 positive cells and gating CD38-V450 positive blasts and negative LSCs. dnFOS transduced/induced PDX were gated for viability on forward/side scatter and sorted for GFP+ as compared to a non-transduced population. dnFOS transduced cell lines were sorted based on forward/side-scatter only to single cells.

#### CyTOF Panel design and in-house labelling of purified antibodies

The AML CyTOF panel was designed to include cell markers specific for myeloid blasts and cell signalling markers of interest. For most of the targets, antibodies were acquired in pre-conjugated format from the Standard BioTools catalogue. For other targets we performed in-house custom conjugations using the MaxPar X8 antibody-labelling kit (Standard BioTools) following the manufacturers protocol. In addition to lanthanide metals, Indium-115 (Sigma Aldrich) and Platinum- 198 (Fluidigm) were used to label antibodies.

Briefly, X8 polymer stored at -20°C was thawed, resuspended in L buffer and then loaded with 50 mM of lanthanide metal (or In115) at 37°C for 40 mins. Metal loaded polymers were washed twice, firstly with L buffer and 25 mins centrifugation, and then with C buffer in a 30 mins centrifugation step. During the polymer wash steps 100 µg of purified antibodies were washed with R buffer using a 50kDa centrifugal unit. Antibodies were then partially reduced with 4 mM TCEP (Fisher) for 30 mins at 37°C. Reduced antibodies were twice washed in C buffer. Partially reduced antibodies were mixed with metal-loaded polymer and incubated at 37°C for 90 mins. Conjugated antibodies were washed and centrifuged four times using W buffer. Purified labelled antibodies were finally eluted from the 50kDa units by a centrifugation step using 100 µL of W buffer and assessed for protein concentration using a NanoDrop spectrophotometer (ThermoFisher). The antibody preparations were returned to the 50kDa units for a final buffer exchange step with 100 µl PBS antibody stabilization buffer (Candor). For Pt198 labelling we followed the Maecker lab protocol^84^ where platinum directly labels the reduced antibody without the use of polymer. All antibodies were tested at different titres to ascertain the optimal final dilution (Table 2).

Primary bone marrow-derived white blood cells were sorted for CD34 positivity using a CD34 MicroBead Kit (Miltenyi Biotec) and cultured for 10 days as detailed above (primary cultures) such that cells were actively proliferating. Cells were taken and resuspended to 20-30×10^6^/ml. Antibody cocktail was prepared in excess and filtered through a 0.1 µm centrifugal filter column (Merck Millipore) to remove antibody aggregates.

Samples were initially barcoded by staining cells with metal labelled CD298/B2M antibodies for 20min at room temperature (RT). Samples were washed twice with MACS buffer. Resuspended cells were then pooled into a single tube and incubated with Tru-Stain Fc blocking solution (Biolegend) for 10 mins at RT. This was immediately followed by incubation with the surface marker antibody cocktail. Staining was performed at RT for 30min with gentle agitation every 10 min. During the last 2min of the 30min incubation, cells were incubated with Cell ID Cisplatin-194 (Pt194). The Pt194 was then quenched with 3mL MACS buffer. Cells were centrifuged and resuspended in freshly prepared 1.6% paraformaldehyde (Thermo Fisher) and incubated in the dark for 15 mins at RT. Cells were washed in MACS buffer then pelleted cells held on ice for 15 mins. After a further gentle agitation to ensure cells were well dispersed, 1mL of cold methanol was added to each tube. Cells were incubated at -20*°*C overnight. The next day tubes were allowed to reach RT then washed twice with MACS buffer. Cells were incubated with antibodies for intracellular targets for 30 mins at RT. Cells were washed with MACS buffer then stained with 500 µM Rh103 DNA intercalator diluted 1:2000 in 500 ul Fix and Perm buffer (Standard BioTools) at 4°C overnight.

Samples were acquired within 72hr of cell staining. Prior to acquisition, the samples were washed once with MACS buffer and then twice with freshly dispensed milliQ deionized distilled water (ddH_2_O). Cells were then resuspended in ddH_2_O containing 1/10 diluted four element (EQ) normalization beads (Standard BioTools) and filtered through a cell strainer cap (Thermo Fisher). Cell densities were corrected to be lower than 1×10^6^ cells/ml. Samples were then acquired on a Helios mass cytometer (Standard BioTools) at flow rate of 30 µl/min using a standardized acquisition template following routine tuning and instrument optimization using the HT Helios injector. To ensure absence of sample carryover to the next sample, tubes with milliQ ddH_2_O (3min), then wash (nitric acid) solution (3min) and again miliQ ddH_2_O (5min) were run on the instrument in between each sample.

Raw fcs datafiles were (EQ-)bead-normalized using the processing tool in the Fluidigm CyTOF acquisition software. Normalized fcs datafiles were then exported and uploaded to Cytobank software (Beckman Coulter). Each file was cleaned up by a series of manually set gates to exclude normalization beads, non-cellular debris, doublets and dead cells. The processed data was exported into a new experiment where debarcoding was performed to generate individual sample fcs files for further analysis. Processed datafiles were analysed using manual gating using CD45/CD34/CD117 to focus on bulk myeloid cells, then further gated for CD38+/- to focus on LSCs or blasts. Mean ion count data for each channel was exported after confirming normal distribution using biaxial plots and visualised using heatmaps in R. FCS files of gated cells were exported and read into FlowCore in R, ion counts were log_2_ transformed and a pseudocount of 1 added, then a Student’s t-test performed.

#### LSC proliferation assay

Blood from patient 2 underwent lymphoprep and the cells were sorted using the strategy above for LSCs and blasts. Each population were divided into two, and the membranes stained with 1) PKH-26 (Merck) and 2) Claret (Merck). The PKH-26 blasts were combined back with the claret LSCs and vice versa, maintaining the original blast:LSC ratio. These cells were then again divided into two and incubated for 6 days in SFEMII media as described above (without hMSCs to avoid contamination), with 20 µM EdU, and with or without 50ng/ml VEGF and 10ng/ml IL-5. After 6 days the cells were stained for EdU with the EdU proliferation kit iFluor 488 (Abcam) and flow cytometry was carried out using a CytoFlex (Beckman Coulter). Cells were gated for viability using forward/side scatter, then LSCs/Blasts using PKH-26 (PE) vs Claret (APC) and finally EdU positive/negative (FITC). Gating for PKH-26 and Claret was set using cells which were stained in a known proportion of 90:10 PKH- 26:Claret and 10:90 PKH-26:Claret.

#### Immunofluorescence

Cells were adhered to microscope slides using a Shandon Cytospin 4 (Thermo Fisher) at 800 rpm for 3 minutes. A border was drawn using a PAP pen and cells were then fixed with 4% formaldehyde for 10 minutes. Permeabilisation was with PBS/0.1% Triton-X100 for 20 minutes, blocking with PBS/0.1% Tween-20/3% BSA for 1 hour. Mouse anti-FLAG antibody was incubated at 1:100 in PBS/0.1 Tween- 20/1% BSA for 1 hour, room temperature. Alexa fluor 594 goat anti-mouse antibody was incubated at 1:200 in PBS/0.1 Tween-20/1% BSA for 1 hour, room temperature. Slides were mounted using ProLong Gold antifade with DAPI (Invitrogen) then imaged using a Zeiss LSM780 confocal microscope, using a Plan Achromat 40x 1.2 NA water immersion objective, Lasos 30 mW Diode 405 nm and Lasos 2mW HeNe 594 laser lines. Images were acquired using Zen black version 2.1 and post-acquisition brightness and contrast adjustment was performed uniformly across the entire image.

#### Generation of t(8;21) PDX

Frozen bone marrow cells from relapsed patient #4 were transplanted either intrahepatically or intrafemorally as shown in Table 3. PDX cells were harvested from leg and hip bone BM by clearing the bones of all tissue, crushing and washing in PBS to releash the BM. Spleen blasts were isolated by passing through a 50 µM cell sieve. Cells were washed and stored frozen in 10%DMSO/90%FBS. Peripheral blood blasts were sampled from the tail vein (< 10% total blood volume/bleed) and analysed by flow cytometry. Leukemia-inducing cell frequency was calculated by intrafemoral secondary transplantion of PDX isolated from NSG bone marrow, with time to endpoint recorded.

**Table 3:** Creation of t(8;21) patient #4 relapse xenograft. *In vivo inhibition of VEGFA and IL5RA in t(8;21) PDX mice*

| Transplant (tx) route | Number of cells/mouse in 20μl | Mouse strain/age | Number of mice engrafted | Latency (weeks) | Sites of engraftment |
| --- | --- | --- | --- | --- | --- |
| intra-hepatic (1.2Gy 24hr prior to tx) | 1x10E6 | MISTRG / 3 days old | 1/5 | 44 | Abdominal tumour |
| intra-femoral | 2x10E6 | NSG / 8-12 weeks old | 2/2 | 21, 30 | Bone marrow, spleen(weight 0.1g), skull, thymus, liver, ascites, ovary, uterus, abdominal tumours, peripheral blood |
| intra-femoral | 2x10E6 | MISTRG / 14 weeks old | 1/1 | 32 | Bone marrow, spleen(weight 0.1g), lung tumour, ovary, uterus, leg tumour, peripheral blood |

Thirteen NSG mice were each transplanted intra-femorally (as above) with 0.6x10^6^ cells from t(8;21) patient #4 secondary transplanted PDX BM. On day 3 after transplant mice were randomised into treatment groups for intra-peritoneal (i.p.) injection (volume<6ml/kg, 29G U100 insulin syringe with needle) of control-vehicle - saline (0.9% NaCl_2_) n=5; Bevacizumab 2mg/kg in saline n=4 and Benralizumab 0.38mg/kg in saline n=4. Dosing was continued twice weekly for 13 doses/mouse. Mice were humanely killed when they reached the endpoints specified above or at 92 days.

#### RNA isolation, cDNA synthesis and qRT-PCR

RNA was isolated from Kasumi-1 cells after 2 days after shRUNX1::ETO knockdown was induced with doxycycline using the Nucleospin RNA kit (Macherey-Nagel). cDNA was synthesised using Superscript II (Invitrogen) from 1 µg total RNA, using oligo(dT)12-18 primer. qRT-PCR was carried out using diluted cDNA, Sybr Green PCR Master Mix (Thermo Fisher), 5 µM of sense and antisense primers (sequences listed in Resource Table).

#### RNA-seq

RNA was isolated using the Nucleospin RNA kit (Macherey-Nagel) for Kasumi-1, or RNeasy Plus micro kit (Qiagen) for patient/PDX cells. RNA libraries were generated using TruSeq stranded total RNA library prep kit with ribo-zero for Kasumi-1, or NEBNext Ultra II Directional RNA Library prep kit (New England Biolabs) for primary cells, per the manufacturer’s instructions. Illumina sequencing was performed on a NextSeq 550 run in paired-end mode for 150 cycles.

#### scRNA-seq

Patient cells were sorted for LSCs and blasts as described above, then re-combined at a 1:1 LSC:blast ratio, with 30000 total cells in 45 µl. Cell viability was confirmed then loaded on a Chromium Single Cell Instrument (10X Genomics), to recover 5000 single cells. Library generation was performed by the Genomics Birmingham sequencing facility using the Chromium single cell 3’ library and gel bead kit v3.1. Illumina sequencing was performed on a NovaSeq 6000 S1 run in paired-end mode for 150 cycles at a depth of 20000 reads per cell.

#### DNaseI-seq

DNaseI digestions were performed as in Bert et al.^85^ Cells were permeabilized in DNaseI resuspension buffer (60mM KCl, 10 mM Tris pH7.4, 15 mM NaCl, 5 mM MgCl2 and 300 mM sucrose) and then DNaseI diluted in dilution buffer (60mM KCl, 0.4% NP40, 15 mM NaCl, 5 mM MgCl2, 10 mM Tris pH 7.4 and 2mM CaCl2 was added and incubated at 22°C for exactly 3 minutes. The digestion was terminated by adding cell lysis buffer (300 mM Sodium Acetate, 10 mM EDTA pH 7.4, 1% SDS and 1 mg/ml proteinase K). DNA was purified using phenol-chloroform extraction. Library preparation was performed using the KAPA HyperPrep kit (Roche) on extracted DNA with size selection for 200-300 bp fragments and sequenced on a NextSeq 550 (Illumina) run in single-end mode for 75 cycles.

#### ATAC-seq

Omni ATAC-seq was performed as in Corces et al.^86^. Briefly, cells were washed in ATAC resuspension buffer (RSB) (10 mM Tris-HCl pH7.5, 10 mM NaCl and 3 mM MgCl2) and then lysed for 3 minutes on ice in RSB buffer with 0.1% NP-40, 0.1% Tween-20. Then the cells were washed with 1 ml of ATAC wash buffer consisting of RSB with 0.1% Tween-20. Nuclei were resuspended in ATAC transposition buffer consisting of 25μl TD buffer and a concentration of Tn5 transposase enzyme (Illumina) related to the number of input cells up to 2.5 μl, 16.5 μl PBS, 5 μl water, 0.1% tween-20 and 0.01% digitonin and then incubated on a thermomixer at 37°C for 30 minutes. The transposed DNA was then amplified by PCR amplification up to ¼ of maximum amplification, as assessed by a qPCR side reaction and sequenced on a NextSeq 550 (Illumina) run in single-end mode for 75 cycles.

#### ChIP-seq

Between 2 and 20 x10^6^ cells (number is antibody dependent) were crosslinked following 72 hours of dnFOS induction with doxycycline, using 1% formaldehyde for 10 minutes at room temperature. For GATA2 and FOS cells were double crosslinked, by adding 415 µg/ml Di(N-succinimidyl) glutarate for 45 minutes prior to formaldehyde crosslinking. Cells were lysed and nuclei extracted using lysis buffer (10 mM HEPES pH 8.0, 10 mM EDTA pH 8.0, 0.5 mM EGTA pH 8.0, 0.25% Triton X-100, protease inhibitor cocktail (PIC) 1:100) followed by nuclear lysis buffer (10 mM HEPES pH 8.0, 1 mM EDTA pH 8.0, 0.5 mM EGTA pH 8.0, 0.01% Triton X-100, 200mM NaCl, PIC 1:100). Nuclei were sheared to around 100-600 bp in sonication buffer (25 mM Tris pH 8.0, 150 mM NaCl, 2 mM EDTA pH 8.0, 1% Triton X-100, 0.25% SDS, PIC 1:100), using a Picoruptor (Diagenode) for between 4 and 16 cycles of 30s on/30s off (cycle number dependent on cell number and crosslinking). Sheared chromatin was diluted in IP buffer (25 mM Tris 1M pH 8.0, 150 mM NaCl, 2 mM EDTA pH 8.0, 1% Triton X-100, 7.5% Glycerol, PIC 1:1000). Dynabeads protein G were pre-incubated with antibodies against FOS, CEBPA, RUNX1, RUNX1::ETO, PU.1 GATA2, H3K27ac, H3K9acS10P or H3K4me3 (all details in resource table) for 2 hours at 4°, then added to the chromatin. Chromatin and antibody-beads mixture were incubated for between 4 and 18 hours (antibody dependent) at 4°. Beads were then washed sequentially: once with buffer 1 (20 mM Tris pH 8.0, 150 mM NaCl, 2 mM EDTA pH 8.0, 1% Triton X-100, 0.1% SDS), twice with buffer 2 (20 mM Tris pH 8.0, 500 mM NaCl, 2 mM EDTA pH 8.0, 1% Triton X-100, 0.1% SDS), once with buffer 3 (10 mM Tris pH 8.0, 250 mM LiCl, EDTA pH 8.0, 0.5% NP-40, 0.5% Sodium deoxycholate), twice with buffer 4 (10 mM Tris pH 8.0, 50 mM NaCl, 1 mM EDTA pH 8.0). Enriched DNA was eluted from the beads with 100 mM sodium bicarbonate and 1% SDS. Crosslinks were reversed with 25 µg Proteinase K for 16 hours at 65°C and DNA was purified using AmpureXP beads (Beckman Coulter). Enrichment was confirmed using qRT-PCR with known positive and negative binding sites for each protein target, then library preparation and sequencing was carried out as for DNaseI-seq with size selection for 200-500 bp fragments.

#### CUT&RUN

Nuclear CUT&RUN was performed as in Skene and Henikoff^87^. Briefly, 1x105 cells were washed with PBS. Nuclei were isolated with NE Buffer (20 mM Hepes-KOH pH 7.9, 10 mM KCl, 0.5 mM spermidine, 0.1% Triton X-100, 20% Glycerol), captured with Concanavalin A beads (Bangs Laboratories, BP531) and incubated with anti-H3K27me3 antibody (Cell signalling, C36B11) for 2 h at 4°C. After washing away unbound antibody with wash buffer (20 mM HEPES-NaOH pH 7.4, 150 mM NaCl, 0.5 mM Spermidine, 0.1% BSA and 1x protease inhibitor cocktails from Sigma), protein A-MNase (provided by the Henikoff laboratory) was added at a 1:200 ratio and incubated for 1 h at 4°C. The nuclei were washed again and were equilibrated to 0°C on a metal block and MNase digestion was activated with CaCl2 at a final concentration of 2mM for 5 minutes. The digestion was terminated with the addition of equal volume of 2xSTOP buffer (200 mM NaCl, 20 mM EDTA, 4 mM EGTA, 50 mg/mL RNase A and 40 mg/mL glycogen). The protein-DNA complex was released by centrifugation and then digested by proteinase K at 70°C for 10 minutes and DNA was purified using phenol-chloroform extraction. Library preparation was performed using the KAPA HyperPrep kit (Roche) on extracted DNA and sequenced on a NextSeq 550 (Illumina) run in single-end mode for 75 cycles.

#### RNA-seq analysis

Raw paired-end reads were processed with Trimmomatic v0.39 to remove sequencing adaptors and low-quality sequences. The processes reads were then aligned to the human genome (version hg38) using Hisat2 v2.2.1 with default parameters.

Gene expression from sorted LSC and blast experiments were calculated as fragments per kilobase of transcript per million mapped reads (FPKM) using Stringtie v2.1.3 with default parameters and gene models from Ensembl as the reference transcriptome. Only protein-coding genes that were expressed with an FPKM value > 1 in either the LSC or blast samples were retained for further analysis. FPKM values were normalized using upper-quartile normalization and further log2- transformed with a pseudocount of 1 added before transformation. A gene was considered to be either LSC or blast specific if it had a fold-change > 1 between cell types.

Counts from all other RNA-Seq experiments were obtained using featureCounts from the Subread package v2.0.1 using the options -p -B -s2 and gene models from refSeq as the reference transcriptome. Only genes with at least 50 counts in at least one sample were retained for further analysis. Counts were normalized using the edgeR package in R v4.1.0, and differential gene expression analysis was then carried out using limma-voom. For experments where replicates were available, a gene was considered to be differentially expressed if it had a fold-change of at least 2 and a Benjamini-Hochberg adjusted p-value < 0.1. In cases where no replicates were possible, only a 2-fold-change was used.

Gene set enrichment analysis (GSEA) was carried out using the GSEA software (Broad Institute). Genes were ranked by the log_2_ fold change and enrichment was calculated for gene sets comprising the LSC or Blast specific differential genes.

Gene ontology (GO) term analysis was carried out using DAVID 6.8. GO terms were ranked according to p-value, with the top 10 statistically significant terms (adj. p-value < 0.005) selected for further analysis. GO term results were then visualised as a bubble-plot in R v4.1.0 with the size of each bubble representing the percentage of genes from that GO term that were present in the set of differentially expressed genes, and the colour corresponding to the adjusted p-value.

#### ATAC/DNaseI-seq analysis

ATAC or DNaseI sequencing reads were processed with Trimmomatic v0.39 to remove sequencing adaptors and low-quality reads. Trimmed reads were aligned to the human genome (version hg38) using Bowtie2 v2.3.5.1 using the setting --very-sensitive-local. PCR duplicates were removed using the MarkDuplicates function in Picard 2.21.1. Bigwig files were made using the bamCoverage function in deepTools 3.5.0 and normalised as counts per million (CPM). These bigwig files were then plotted using the UCSC genome browser. Peaks were called using MACS2 v2.2.7.1 using the settings -q 0.0005 -B --trackline --nomodel --shift -100 --extsize 200.

To carry out differential chromatin accessibility analysis, a peak union was generated using the bedtools v2.29.2 merge function. The average tag-density in a 400-bp window centred on the peak union summits was calculated for each sample using the annotatePeaks.pl function in Homer v4.11 using the bedGraph files generated by MACS2. These were then normalised as CPM and further log2-transformed as log2(CPM + 0.1). Peaks were considered to be differentially accessible if there was at least a 2-fold difference between samples.

Density plots were generated using Homer v4.11 annotatePeaks.pl function using the bedGraph files generated by MACS2, with the options -size 2000 -hist 10 -ghist and plotted using JavaTreeView 1.1.6.

In order to measure if a transcription factor motif was overrepresented in a set of differentially accessible peaks, we calculated a motif enrichment score (ES) as follows. The number of motifs in a peak set was first counted by extracting the motif positions using the findMotifsGenome.pl function in Homer with the options -size 200 -find. The probability weight matrices provided by the Homer motif database were used in all analyses. The enrichment score was then calculated as:

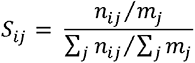

where i is the motif, j is the peak set, n_ij_ is the number of sites in peak set j that contain the motif i and m_j_ is the total number of sites in peak set j. The scores were then hierarchically clustered using complete linkage of the Euclidean distance in R and displayed as a heat map.

Average profiles were created using normalized bigwig files. To do this, the average peak height for each sample was calculated for each sample using the computeMatrix and plotProfile functions in deepTools. A normalization factor was then calculated for each sample so that the average peak height was the same for all samples. Normalized bigwig files were then created using the bamCoverage function in deepTools using the --scale option to apply the normalization factor. The average profile was then plotted using the computeMatrix and plotProfile functions.

Average motif profiles were generated using Homer annotatePeaks.pl with the options -size 2000 - hist 10 -m <target motif position weight matrices> and plotted using R ggplot2 using the geom_smooth loess function.

#### ChIP-seq/CUT&RUN analysis

ChIP-sequencing reads were processed with Trimmomatic v0.39 to remove sequencing adaptors and low quality reads. Trimmed reads were aligned to the human genome (version hg38) using Bowtie2 v2.3.5.1 using the setting --very-sensitive-local. PCR duplicates were removed using the MarkDuplicates function Picard v2.21.1. Bigwig files were created for viewing in UCSC genome browser using deepTools 3.5.0 bamCoverage, with normalisation using counts per million (CPM). Peaks were called using MACS2 using the settings -q 0.01 -B --trackline. Differential peaks were calculated as for ATAC-seq.

Average profiles were generated as for ATAC-seq, except for H3K27me3 where normalisation was only by counts per million due to the broad regions which have this mark. The average peak height was calculated from these profiles at specific sites and a log_2_ fold change calculated and plotted as a heatmap in R using hierarchical clustering as for ES above. ES and density plots were generated as for ATAC-seq except for H3K27me3 and H3K4me3 where density plots were generated using deepTools plotHeatmap in conjunction with the average profiles.

In order to ensure that ChIP peaks were associated with the correct target gene we used processed promoter-capture Hi-C data from Assi et al.^7^. This was done by first searching for peaks that could be assigned to a DNaseI hypersensitive site (DHS) for which the Hi-C data could associate the DHS with the correct gene promoter. In cases where no Hi-C association was available, peaks were assigned to their closest gene based on transcription start site (TSS) using the annotatePeaks.pl function in Homer.

#### scRNA-seq analysis

Reads from single-cell RNA-Seq experiments were aligned to the human genome (version hg38) and quantified using the count function in CellRanger v3.1.0 from 10x Genomics and using gene models from Ensembl as the reference transcriptome. Unique Molecular Identifier (UMI) count data was filtered for low quality cells by removing cells with less than 500 and more than 5000 detectable genes. Cells that had more than 20% of UMIs aligned to mitochondrial transcripts were also excluded from further analysis. UMI counts were normalized using the log-normalize method in the Seurat package v4.1.0 in R v4.1.2. The cell cycle stage was then estimated for each cell using the CellCycleScoring function in Seurat and using the in-built lists of cell cycle stage associated genes. To account for the possible effect of cell cycle stage on downstream clustering analysis, S-phase and G2M-phase scores were included as variables in a linear regression model using the ScaleData function in Seurat. Principal Components Analysis (PCA) was then performed on the normalized and scaled data, with the first 20 principal components for t(8;21) #1 and 14 principal components for t(8;21) #2 selected for further analysis. Cells were then clustered using the FindClusters function in Seurat with a resolution value of 0.8 and visualized using Uniform Manifold Approximation and Projection (UMAP). Cluster marker genes, corresponding to genes that are significantly higher expressed in a cluster compared to all other cells outside of that cluster, were identified using the FindAllMarkers function. Genes that had an average log2-fold change of at least 0.25 with an adjusted p-value less than 0.1 were selected as marker genes.

In order to classify a single-cell cluster as either blast or LSC, specific genes from the blast and LSC bulk RNA-seq above were used as a reference gene expression signature for Gene Set Enrichment Analysis (GSEA). GSEA was carried out using the fgsea package v1.10.1 (27) in R. To do this, cluster marker genes from single-cell clusters were used as pathways and compared to the gene expression signatures derived from the bulk data. This analysis produced a normalised enrichment score (NES) for each cluster, with a positive NES suggesting that a cluster has a more blast-like gene expression signature and a negative NES suggesting a more LSC-like signature. Only clusters with a Benjamini-Hochberg adjusted p-value < 0.05 and an absolute NES >1 were considered to be positively classified as either LSC or blast.

Single-cell trajectory analysis was carried out using Monocle3 v1.0.0^82^. Processed data from Seurat was imported to Monocle and trajectories were inferred using the learn_graph function. Pseudotime was then calculated using the order_cells command, using cells from the earliest inferred LSC population as the root. Trajectories were then plotted on the UMAP calculated by Seurat.

Z-scores of t(8;21)-specific genes were calculated by first calculating the average gene expression per cluster using the AverageExpression function in Seurat. The t(8;21)-specific genes were calculated using normalised FPKM values from bulk AML samples obtained from Assi et al^7^, with genes considered as t(8;21)-specific if they were at least 2-fold higher in the average of all t(8;21) patients compared to the average of each of the other AML subtypes or PBSCs. The average cluster expression of the t(8;21)-specific set of genes was then transformed to a Z-score using the scale function in R and plotted as a heatmap with supervised clustering by cell cluster ordered by the inferred pseudotime trajectory and ordered from highest to lowest Z score in each population.

Genes that were specifically differential in G_0_/G_1_ cells were obtained by subsetting all of the non-S/G2M phase cells based on the cell cycle scoring above. The FindAllMarkers function was then run on this subset using the LSC/Blast classification rather than the clusters. Genes that had an average log2-fold change of at least 0.25 with an adjusted p-value less than 0.05 were selected as marker genes for LSCs or blasts.

#### Quantification and statistical analysis

For comparisons of drug/cytokine treatment vs control only two-sided Student’s t-tests were performed. For growth curves two-way ANOVA was performed with Dunnett correction for multiple comparisons at each time point. For mass cytometry data Student’s t-tests were performed on log_2_ transformed data.

## Supplemental Table Titles

Supplemental Table1 – 2-fold LSC and blast specific genes from primary AML patient RNA-seq datasets

Supplemental Table 2 - G1-stage LSC and Blast marker genes and associated GO terms from primary AML patient scRNA-seq

Supplemental Table 3 - RNA fold changes following expression of dnFOS in Kasumi-1

Supplemental Table 4 - RNA fold changes following expression of dnFOS in PDX or healthy CD34+ cells

## Notes

### Competing Interest Statement

The authors have declared no competing interest.

### Summary of Updates

Addition of experiments using IL5RA inhibitor to Figures 3 and 4

