## Supplemental Figures for "Leukemic stem cells hijack lineage inappropriate signalling pathways to promote their growth"

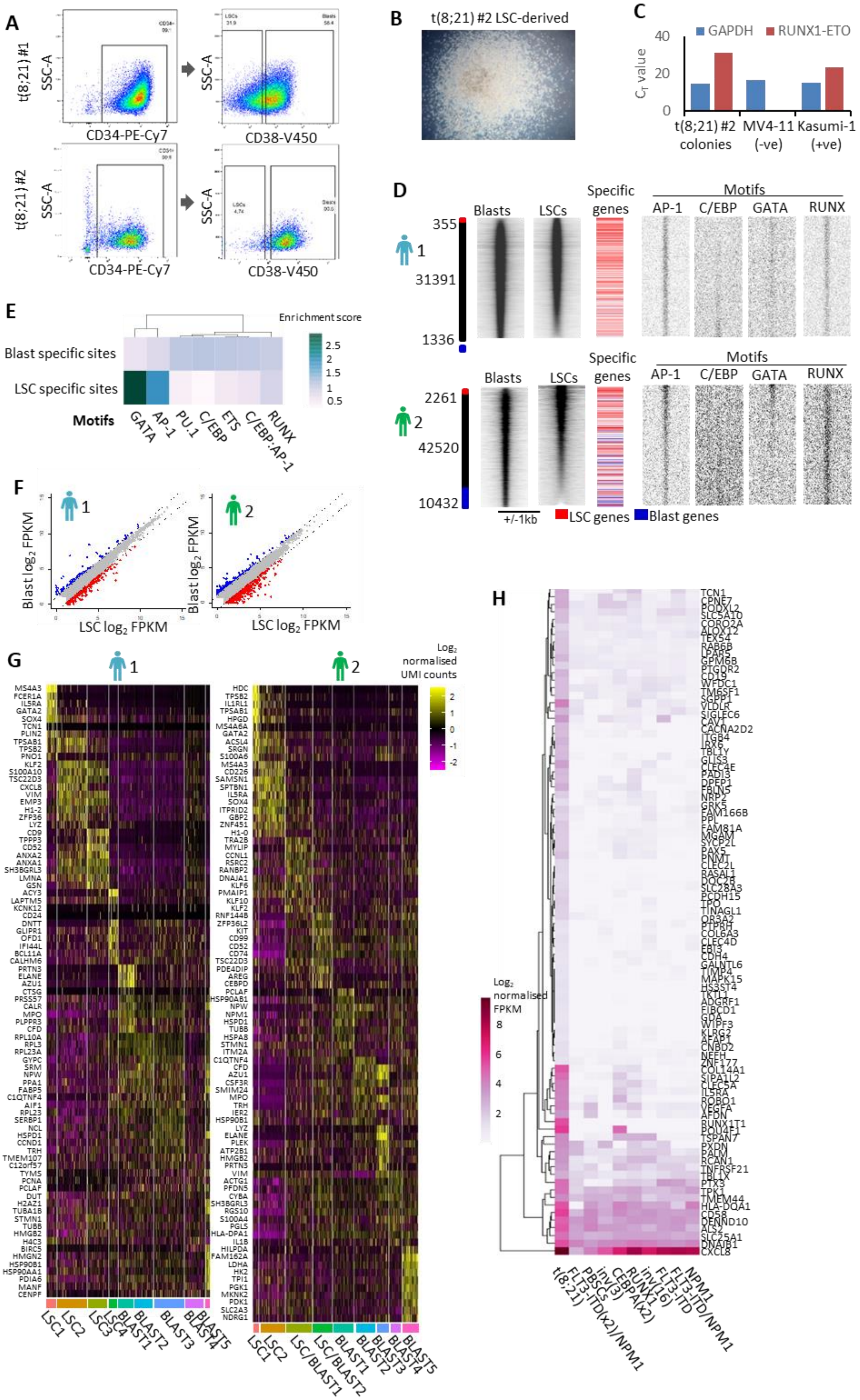

A
t(8;21) #1
t(8;21) #2
SSC-A
CD34-PE-Cy7
CD38-V450
B
t(8;21) #2 LSC-derived
C
GAPDH
RUNX1-ETO
40
20
0
t(8;21) #2 colonies
MV4-11 (-ve)
Kasumi-1 (+ve)
D
Blasts
LSCs
Specific genes
Motifs
AP-1
C/EBP
GATA
RUNX
355
1
31391
1336
2261
2
42520
10432
+/-1kb
LSC genes
Blast genes
E
Enrichment score
2.5
2
1.5
1
0.5
Blast specific sites
LSC specific sites
Motifs
GATA
AP-1
PU.1
C/EBP
ETS
C/EBP:AP-1
RUNX
F
1
2
Blast log2 FPKM
LSC log2 FPKM
G
1
2
Log2 normalised UMI counts
2
1
0
-1
-2
LSC1
LSC2
LSC3
LSC4
BLAST1
BLAST2
BLAST3
BLAST4
BLAST5
LSC/BLAST1
LSC/BLAST2
H
Log2 normalised FPKM
8
6
4
2
t(8;21)
FLT3-ITD(x2)/NPM1
PBSCs
inv(3)
CEBPA(x2)
RUNX1
inv(16)
FLT3-ITD
FLT3-ITD/NPM1
NPM1

**Supplemental Figure 1: Subtype specific gene expression and chromatin accessibility is established in LSCs**

**(A)** Biplot showing the population of cells sorted for CD34 and CD38 for the experiments in Figure 1. **(B)** Representative light microscope image of one of the colonies formed from t(8;21) #2 LSCs, no colonies were observed from the blasts. **(C)** qRT-PCR was performed the colonies in B, RUNX1::ETO was detected with primers which span the breakpoint albeit at a lower expression level than Kasumi-1 cell line. **(D)** ATAC-seq on sorted LSCs and blasts was ranked by fold change of the tag count at distal peaks and represented as density plots (+/1kb of the summit). The blue bar indicates blast specific sites and the red bar LSC specific sites where the normalised tag-count of specific sites is at least two-fold different. Transcription factor binding motif frequencies are plotted along the same axis across the same window, and t(8;21) LSC and blast specific genes are plotted alongside their associated sites in red or blue respectively. **(E)** A motif enrichment score was calculated based on motif frequency in the LSC and blast specific sites calculated from the merged ATAC-seq data from patient #1 and #2. **(F)** Scatter graphs showing the $log_2$ FPKM in bulk sorted LSCs vs blasts in both patients. The blue coloured dots indicate genes which are at least 2-fold higher in the blast population and the red dots those which are 2-fold higher in the LSC population. **(G)** Heatmap showing the top 10 most upregulated marker genes identified in each scRNA-seq cluster. **(H)** Heatmap showing the $log_2$ normalised FPKM of t(8;21) specific genes in AML with different driver mutations and healthy CD34+ PBSCs (Assi et al., 2019).

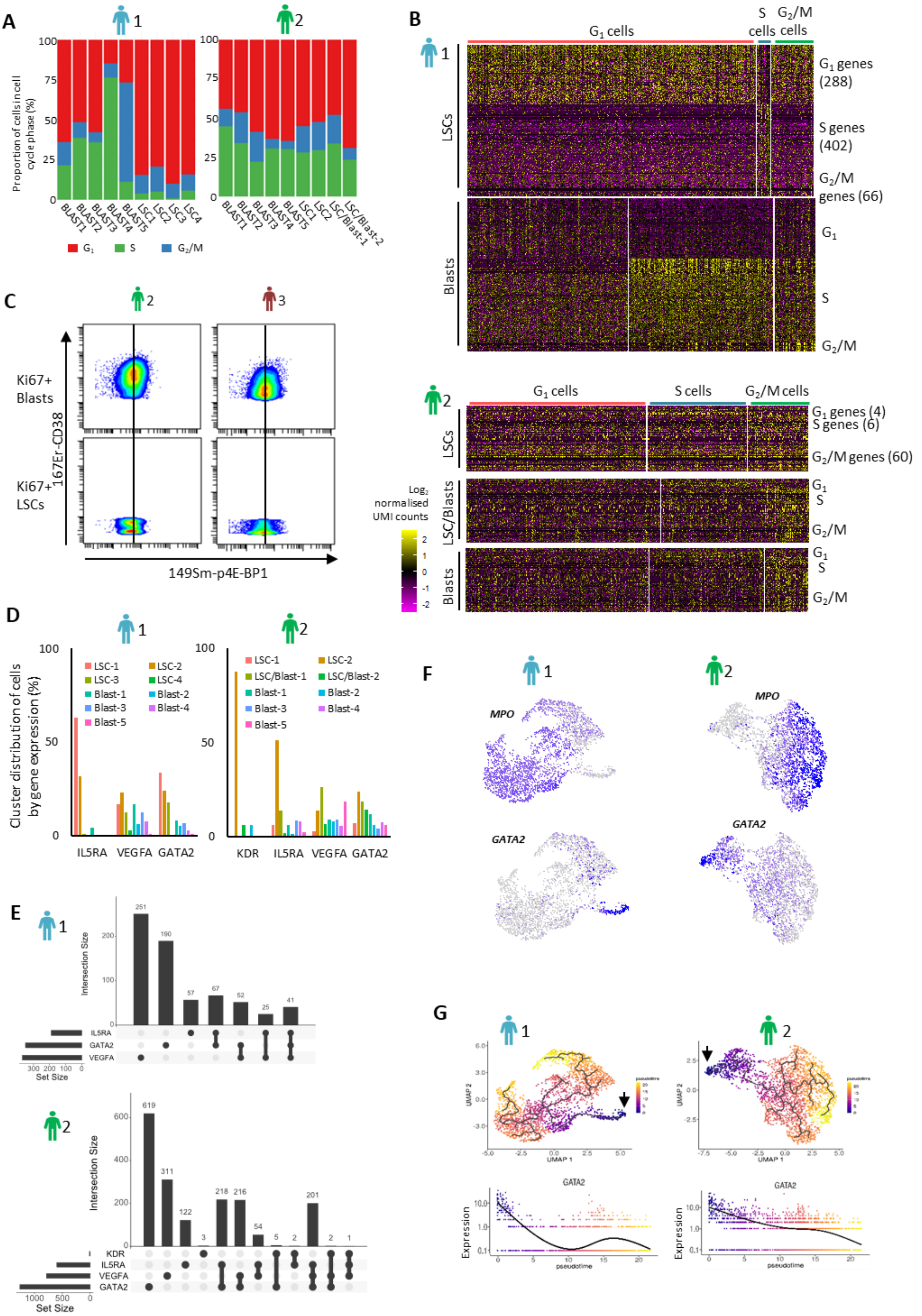
A
1
2
Proportion of cells in cell cycle phase (%)
100
75
50
25
0
BLAST1
BLAST2
BLAST3
BLAST4
BLAST5
LSC1
LSC2
LSC3
LSC4
LSC/Blast-1
LSC/Blast-2
G1
S
G2/M
B
G1 cells
S cells
G2/M cells
LSCs
Blasts
G1 genes (288)
S genes (402)
G2/M genes (66)
G1
S
G2/M
G1 genes (4)
S genes (6)
G2/M genes (60)
LSC/Blasts
Log2 normalised UMI counts
2
1
0
-1
-2
C
3
Ki67+ Blasts
Ki67+ LSCs
167Er-CD38
149Sm-p4E-BP1
D
Cluster distribution of cells by gene expression (%)
LSC-1
LSC-2
LSC-3
LSC-4
Blast-1
Blast-2
Blast-3
Blast-4
Blast-5
LSC/Blast-1
LSC/Blast-2
IL5RA
VEGFA
GATA2
KDR
E
Intersection Size
Set Size
251
190
57
67
52
25
41
619
311
122
3
218
216
54
5
2
201
2
1
F
MPO
GATA2
G
UMAP 1
UMAP 2
pseudotime
Expression
GATA2

**Supplemental Figure 2: t(8;21) AML LSCs are differentially signalling responsive**

**(A)** The proportion of cells in each cluster assigned to each cell cycle stage by their gene expression pattern. **(B)** Heatmaps showing the expression in scRNA-seq of marker genes for each cell cycle phase, plotted according to the cells assigned to each cell cycle phase in LSCs, blasts and for patient 2 the transition cells. **(C)** Biplots showing mass cytometry data for CD38 vs p4-EBP1 in Ki67+ CD34+/CD38+ blasts (top) and Ki67+ CD34+/CD38- LSCs (bottom), the vertical line indicates the median in the blasts. **(D)** Bar charts indicating to which population the cells expressing the genes belong. **(E)** Upset plots showing the number of cells co-expressing *VEGFA, IL5RA, GATA2* (and *KDR* in patient 2 only). **(F)** Expression of *GATA2* and *MPO* projected onto the UMAP plot, where blue indicates the normalised UMI count. **(G)** Pseudo-time trajectory analysis projected onto the UMAP plots, black arrows indicate the beginning of the trajectories. *GATA2* expression across the trajectory is plotted below.

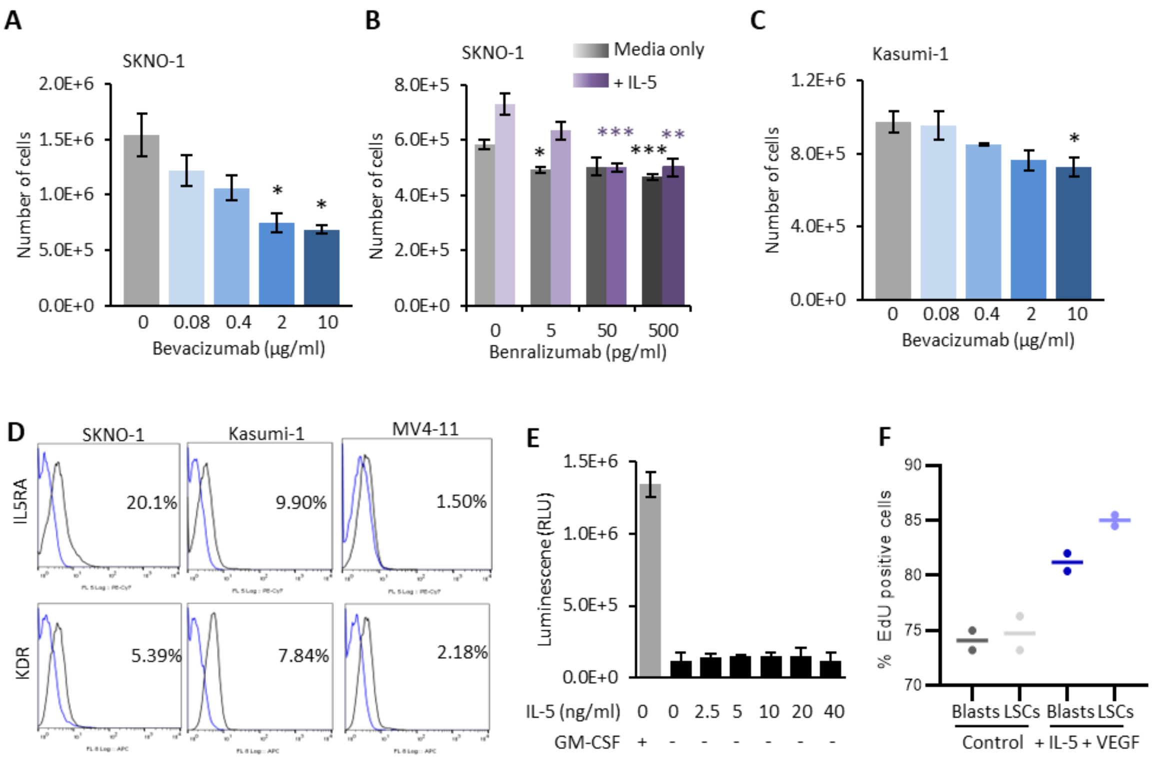


**Supplemental Figure 3: Aberrant VEGF and IL-5 signalling in t(8;21) AML drives LSC activation**

**(A & C)** Dose response showing the number of SKNO-1 **(A)** and Kasumi-1 **(C)** cells after 6 days of treatment with four doses of bevacizumab or with no bevacizumab. **(B)** Dose response showing the number of SKNO-1 cells after 6 days of treatment with benralizumab, +/- IL-5. **(D)** Cell surface expression of IL5RA and KDR was measured by flow cytometry in SKNO-1, Kasumi-1 and MV4-11 cells, representative histograms with isotype control from 4 experiments are shown. **(E)** Dose response showing the luminescence (in relative luminescence units) as determined by celltiter glo at 6 concentrations of IL-5 in the absence of GM-CSF. **(F)** Percentage EdU positivity of PKH-26-LSCs/Claret-Blasts or PKH-26-Blasts/Claret-LSCs, re-combined and cultured for 6 days with or without IL-5 and VEGF. Horizontal bar indicates mean. **(A-C)** * indicates p <0.05, ** p<0.01, *** p<0.005 using Student's T-tests

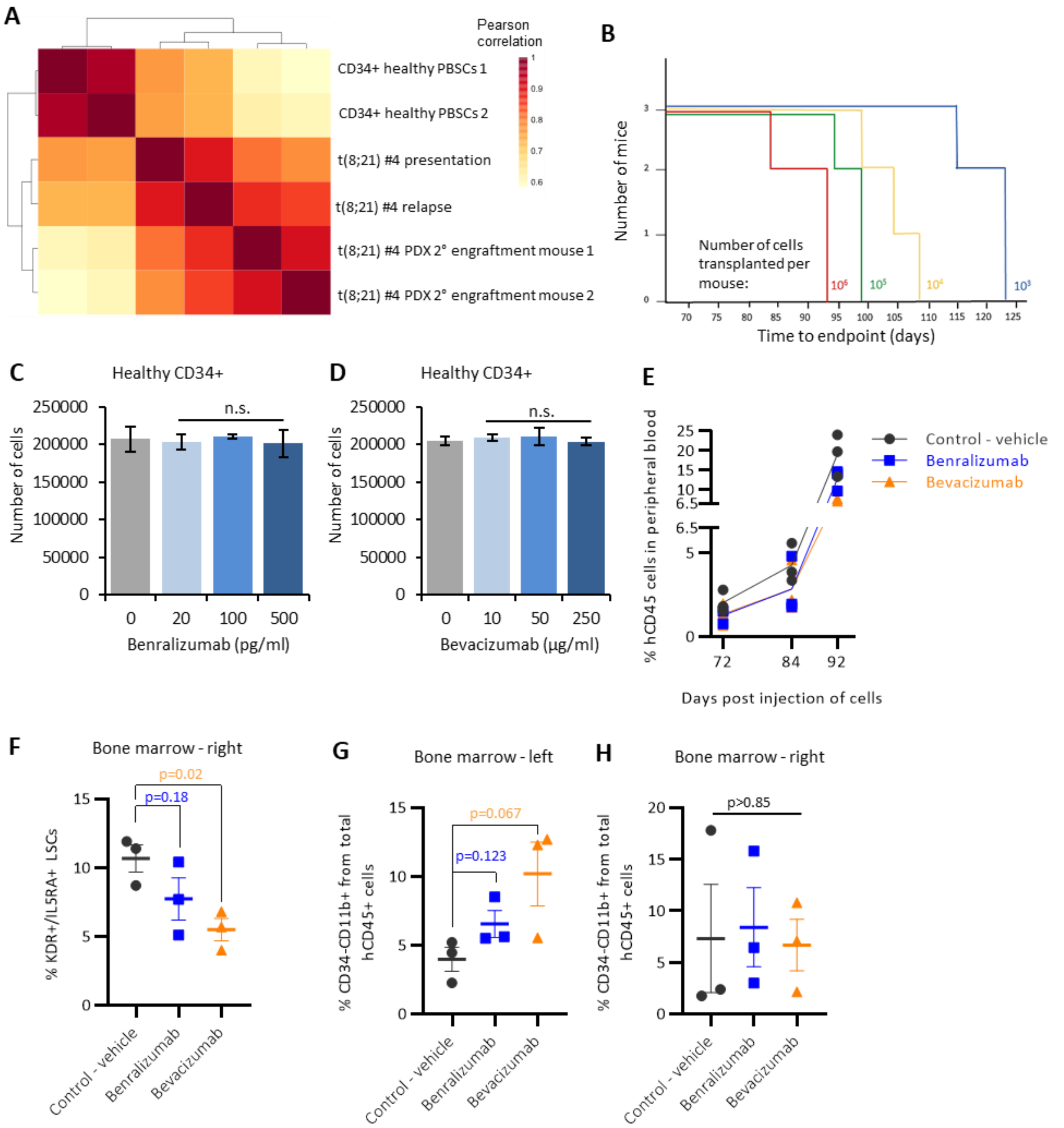


**Supplemental Figure 4: VEGFA and IL5RA inhibitors reduce patient-derived xenograft proliferation**

**(A)** Heatmap with hierarchical clustering showing the Pearson correlation values between gene expression data from RNA-seq on two healthy CD34+ PBSC samples, CD34+ sorted patient cells from t(8;21) #4 at presentation and on relapse, and from two mice which were secondarily engrafted with t(8;21) #4 PDX **(B)** Mice were engrafted with primary PDX cells at $10^3$, $10^4$, $10^5$ and $10^6$ cells per mouse and the survival time until humane end-point recorded **(C-D)** Dose response showing the number of healthy CD34+ cells after 6 days of treatment with 3 doses of benralizumab (+10ng/ml IL-5) **(C)** or with 3 doses of bevacizumab **(D),** bar height shows the mean of 3 replicates and error bars indicate SEM. **(E)** Percentage of human CD45 positive cells in peripheral blood at days 72, 84 and 92 post-injection, p=0.032 by two-way ANOVA for treatment groups. **(F)** Percentage of KDR and IL5RA positive hCD45+/CD34+/CD38- LSCs in right bone marrow at day 92 post-injection. **(G-H)** Percentage of hCD45+/CD34-/CD11b positive cells in left **(G)** and right **(H)** bone marrow at day 92 post-injection. **(F-H)** Horizontal and error bars show mean and SEM of the 3 mice in each treatment group.

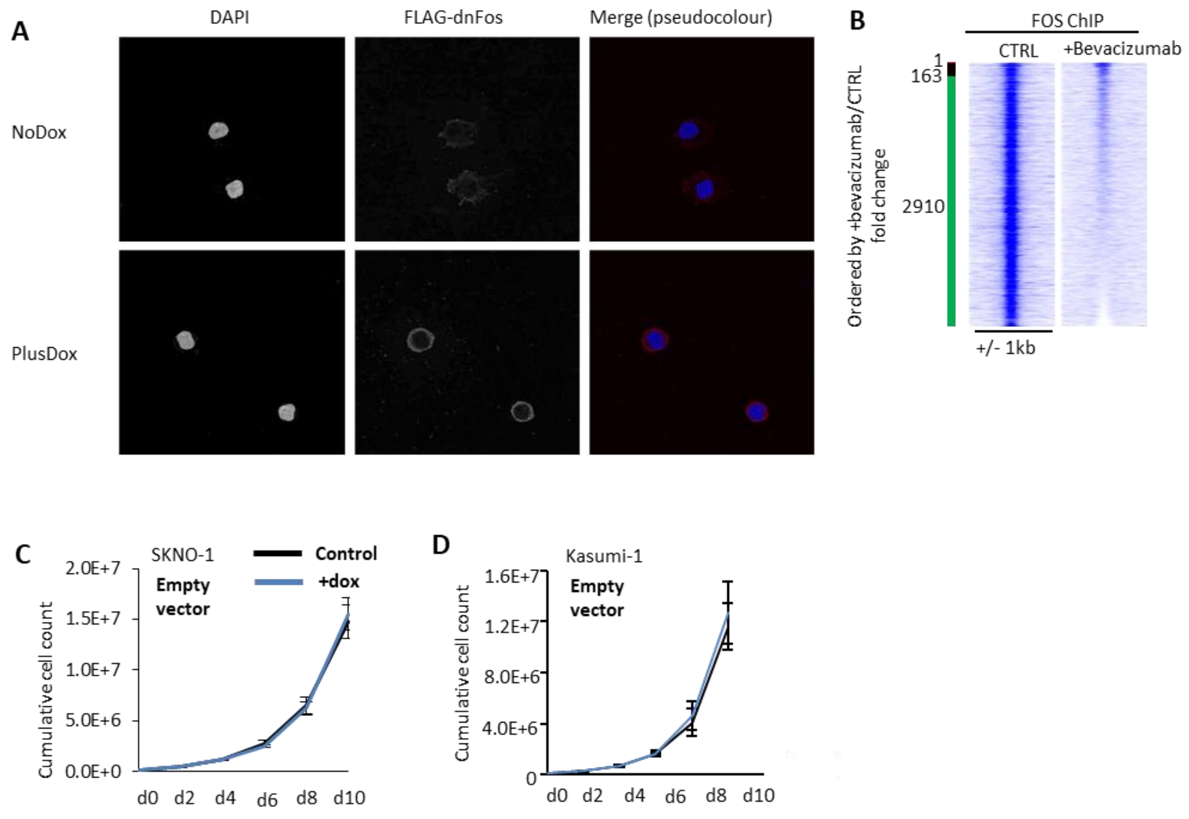


**Supplemental Figure 5: VEGF and IL-5 signals terminate at the AP-1 family of transcription factors**

**(A)** Immunofluorescence images of dnFOS induction in Kasumi-1 cells by staining for the flag-tag, counterstained for DAPI. Separated colour channels are shown in grey scale whilst the pseudocolour is both colour channels combined. **(B)** Density plot showing FOS ChIP signal (+/-1kb from the peak summits) at all sites ranked by the fold change of the tag counts between treated with bevacizumab and an untreated control. The green bar indicates sites which were at least 2-fold reduced with bevacizumab and the red bar indicates the site which was at least 2-fold higher with bevacizumab. **(C-D)** Growth curves were performed by growing SKNO-1 **(C)** or Kasumi-1 **(D)** cells for 10 days, counting and passaging every 2 days following transduction and selection with an EV control of the dnFOS plasmid, where GFP only is induced by doxycycline. Each point indicates the mean of three experiments, error bars show SEM, no significant differences were seen at any time point.

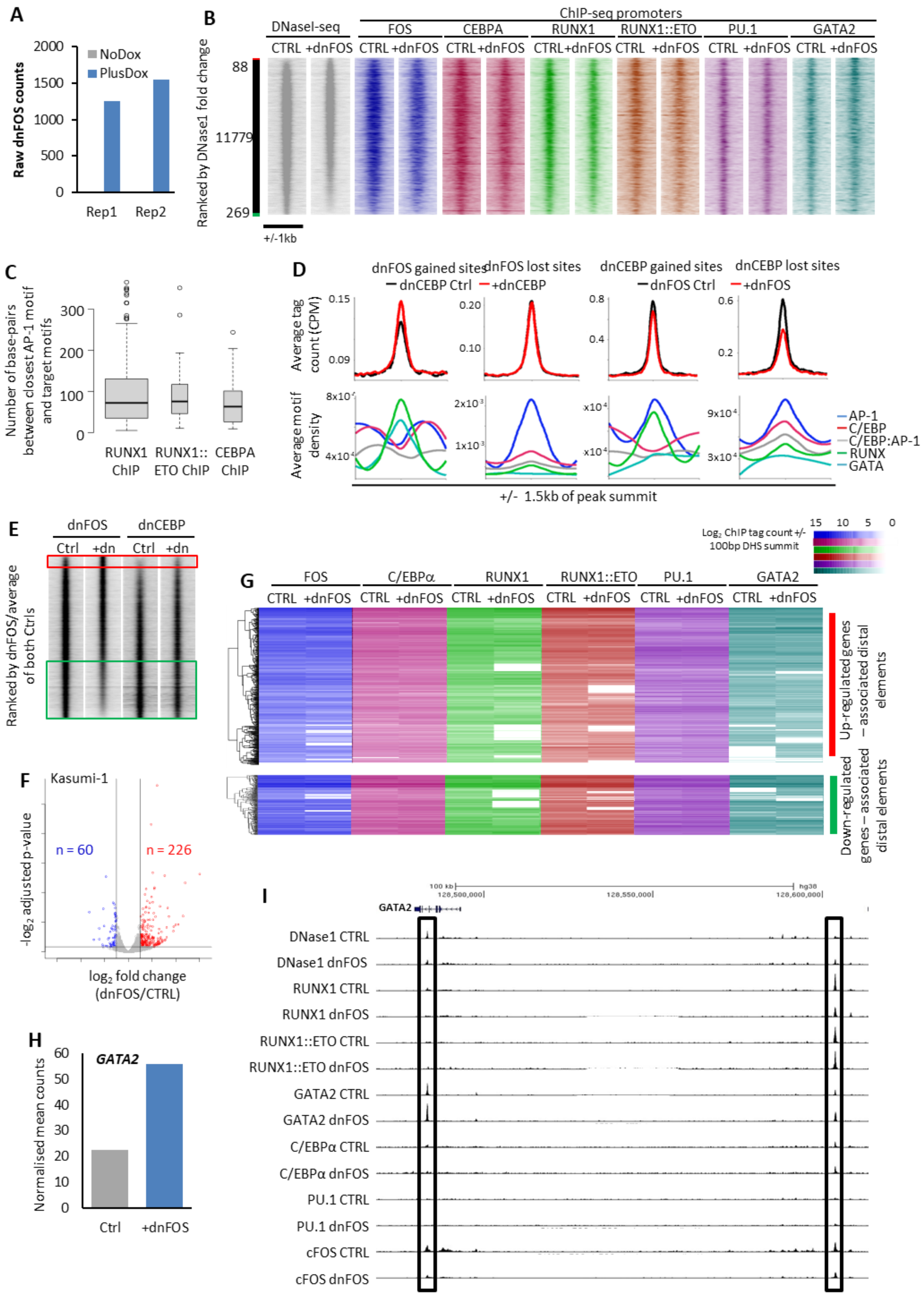


**Supplemental Figure 6: AP-1 orchestrates a shift in transcriptional regulation from an LSC to a blast pattern**

**(A)** Bar chart showing the raw dnFOS counts in the two RNA-seq replicates. **(B**) DNase1 with and without dnFOS induced by doxycycline in the Kasumi-1 cell line, was ranked by fold change of the tag count at promoters and represented as density plots (+/1kb of the summit). The red bar indicates dnFOS specific sites and the green bar control specific sites where the normalised tag-count of specific sites is at least two-fold different. ChIP data from FOS, CEBPA, RUNX1, RUNX1::ETO, PU.1 and GATA2 with and without dnFOS were plotted on the same axis across the same window. **(C)** The proximity of AP-1 motifs to RUNX motifs in RUNX and RUNX1::ETO ChIP control specific sites, or to CEBP motifs in CEBPA ChIP control specific sites was calculated and presented as boxplots where the bold line indicates the mean and the error bars indicate 95% confidence intervals. GSEA analysis was used to compare blast and LSC specific genes which were at least 2-fold differentially expressed in bulk RNA-seq with the ranked fold change gene expression from Kasumi-1 with induction of dominant negative CEBP. **(D)** Average profiles showing the normalised tag count of dnFOS DNaseI at dnCEBP 2-fold specific sites or dnCEBP DNaseI at dnFOS 2-fold specific sites where black shows the controls and red the induced (top) and the average motif density with loess regression at these same sites (bottom). **(E)** Density plots were generated from DNaseI tag count at distal sites, ranked by the fold change of +dnFOS/the average of the dnFOS and dnCEBP controls, +/-1kb of the peak summit, +dnCEBP was plotted on the same axis. **(F)** Volcano plot showing gene expression changes in Kasumi-1 in response to dnFOS induction, vertical lines indicated at least 2-fold change whilst the horizontal line indicates –$log_2$ adjusted p-value of 0.1, genes which are significantly up-regulated are coloured red, and genes which are significantly down-regulated are in blue. **(G)** The tag count of the ChIPs shown in A was measured at all DHS associated with two-fold differentially expressed genes following dnFOS induction (+/- 100bp of the DHS summit). The tag counts and gene expression fold change are shown as heatmaps, with the DHS and their associated gene expression change subject to hierarchical clustering. **(H)** Bar chart showing the normalised mean counts for *GATA2* expression in Kasumi-1 with and without dnFOS. **(I)** UCSC genome browser screenshot showing DNase1 and ChIP-seq at the *GATA2* locus.

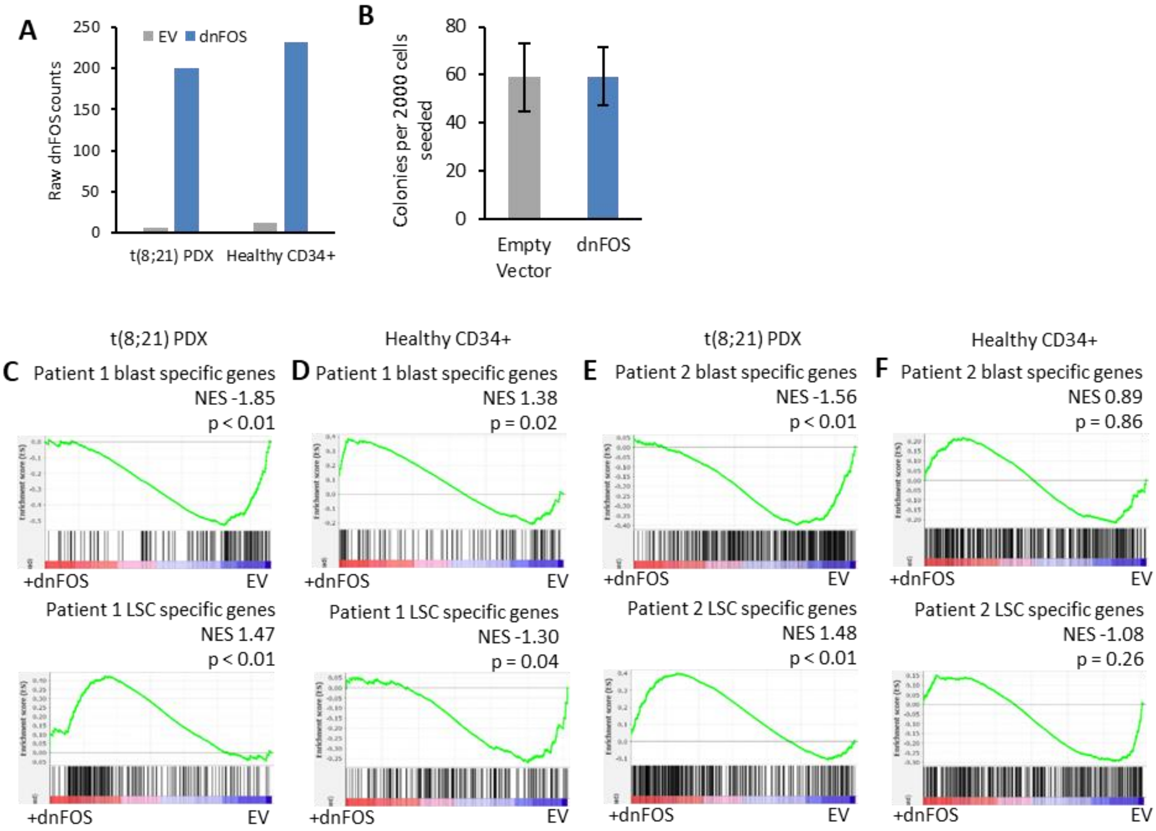


**Supplemental Figure 7: AP-1 is required for maintenance of the blast program**

**(A)** Bar graphs showing the raw number of dnFOS counts in RNA-seq in PDX and healthy CD34+ cells with or without induction of dnFOS. **(B)** Colony forming assay in healthy CD34+ PBSCs induced and sorted for dnFOS or EV. **(C-F)** GSEA was used to compare each individual patients' LSC and blast specific genes with the ranked fold change gene expression from the PDX **(C,E)** and healthy CD34+ cells **(D,F)**, comparing dnFOS to EV. NES shows the normalised enrichment score from the GSEA and the adjusted p-value.

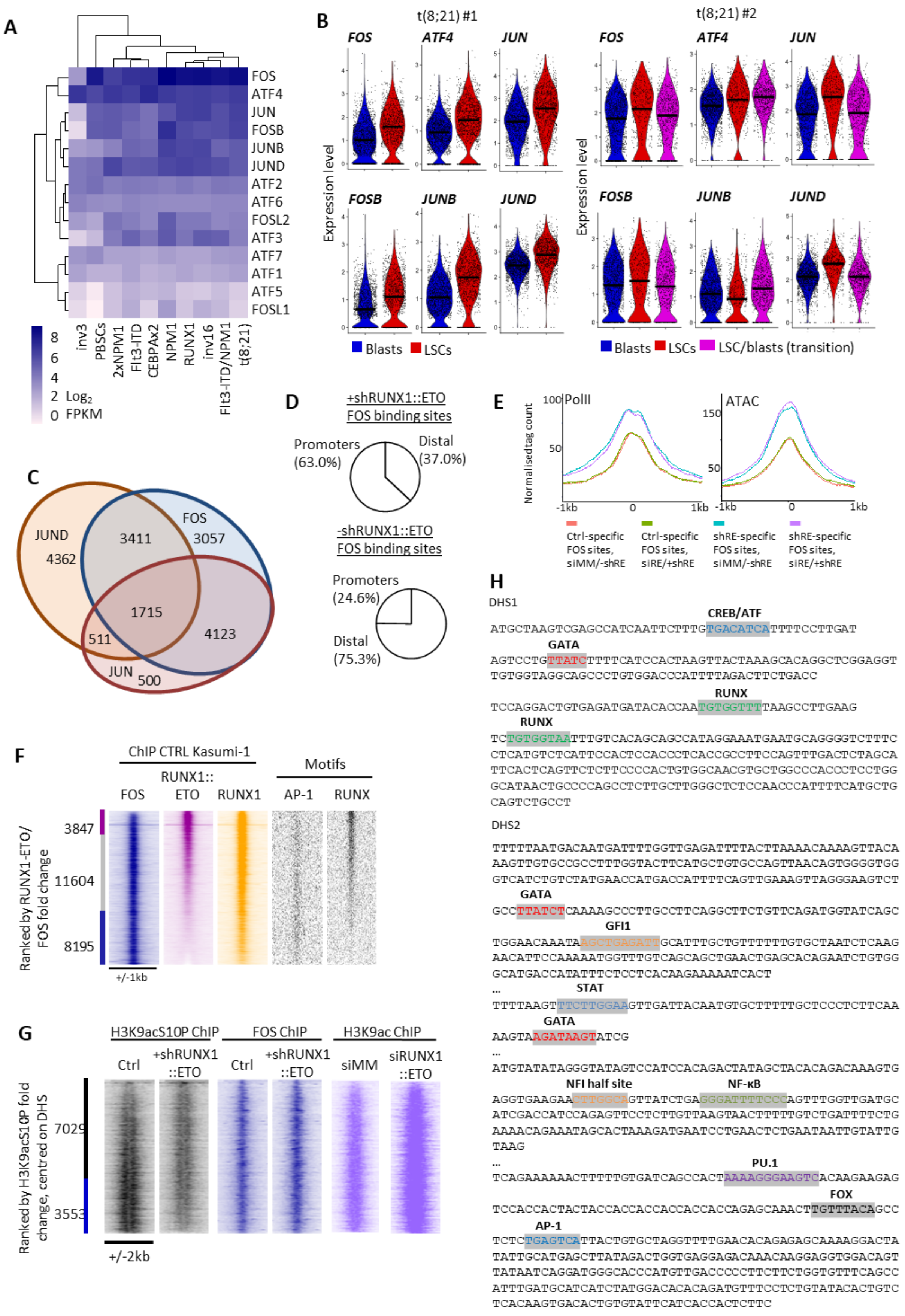
A
FOS
ATF4
JUN
FOSB
JUNB
JUND
ATF2
ATF6
FOSL2
ATF3
ATF7
ATF1
ATF5
FOSL1
8
6
4
2 Log2
0 FPKM
inv3
PBSCs
2xNPM1
Flt3-ITD
CEBPAx2
NPM1
RUNX1
inv16
Flt3-ITD/NPM1
t(8;21)
B
t(8;21) #1
t(8;21) #2
FOS
ATF4
JUN
FOSB
JUNB
JUND
Expression level
Blasts
LSCs
LSC/blasts (transition)
C
JUND
4362
3411
FOS
3057
1715
511
4123
JUN
500
D
+shRUNX1::ETO
FOS binding sites
Promoters
(63.0%)
Distal
(37.0%)
-shRUNX1::ETO
FOS binding sites
Promoters
(24.6%)
Distal
(75.3%)
E
PolII
ATAC
Normalised tag count
-1kb
0
1kb
Ctrl-specific FOS sites, siMM/-shRE
Ctrl-specific FOS sites, siRE/+shRE
shRE-specific FOS sites, siMM/-shRE
shRE-specific FOS sites, siRE/+shRE
F
ChIP CTRL Kasumi-1
Motifs
FOS
RUNX1::ETO
RUNX1
AP-1
RUNX
3847
11604
8195
+/-1kb
Ranked by RUNX1-ETO/FOS fold change
G
H3K9acS10P ChIP
FOS ChIP
H3K9ac ChIP
Ctrl
+shRUNX1::ETO
Ctrl
+shRUNX1::ETO
siMM
siRUNX1::ETO
7029
3553
+/-2kb
Ranked by H3K9acS10P fold change, centred on DHS
H
DHS1
CREB/ATF
GATA
RUNX
RUNX
DHS2
GATA
GFI1
STAT
GATA
NFI half site
NF-κB
PU.1
FOX
AP-1

**Supplemental Figure 8: The signalling response of t(8;21) cells operates within a RUNX1::ETO dependent regulatory circuit**

**(A)** Heatmap with hierarchical clustering showing the average $\log_2$ FPKM of FOS, JUN and ATF transcription factors in 9 subtypes of AML and healthy PBSCs, data from Assi et al. 2019. **(B)** Violin plots showing the expression per cell of the six AP-1 family members most highly expressed in t(8;21) AML, grouped by LSCs and blasts for patient 1 and by LSC, blasts and LSC/blast transition cells for patient 2. The horizontal black bar indicates the median per cell population. **(C)** Venn diagram showing the overlap in ChIP peaks from FOS, JUN and JUND ChIP-seq in Kasumi-1. **(D)** Pie charts showing the proportion of promoters vs distal sites in the specific sites for the FOS ChIP shown in Figure 5B. **(E)** Average profiles showing the tag count, scaled to average peak height, of PolII with siMM or siRUNX1::ETO (Ptasinska et al., 2014) and ATAC-seq with or without doxycycline induction of shRUNX1::ETO at the FOS sites gained and lost with shRUNX1::ETO shown in Figure 5B. **(F)** ChIP for FOS and RUNX1::ETO in Kasumi-1 were ranked by fold change and represented as a density plot. RUNX1 ChIP, AP-1 and RUNX motifs were plotted on the same axis, +/-1kb of the summit. The purple bar indicates RUNX1::ETO specific sites and the blue bar FOS specific sites where the normalised tag-count is at least two-fold different. **(G)** ChIP for H3K9acS10P (note: antibody may also cross-react with H3K9acS28P) was performed with and without shRUNX1::ETO induced by doxycycline in the Kasumi-1 cell line, ranked by fold change of the tag count at promoters and represented as density plots (+/2kb of the summit). The red bar indicates shRUNX1::ETO specific sites and the green bar control specific sites where the normalised tag-count of specific sites is at least two-fold different. FOS ChIP (from Figure 5B) and H3K9ac +/- siRUNX1::ETO (Ptasinska et al., 2012) are plotted along the same axis across the same window. **(H)** Sequences underlying the t(8;21) specific peaks at the *IL5RA* locus with the motif sequences highlighted.
